# Accessory subunits of PRC2 mimic H3K27me3 to restrict the spread of Polycomb domains

**DOI:** 10.1101/2025.03.13.642955

**Authors:** Samuel C Agius, Marion Boudes, Evan Healy, Yoona Kim, Yi-Cheng Chang, Melanie Murray, Jan Manent, Jeanette Rientjes, Zhihao Lai, Sophie Nixon, Jihane Homman-Ludiye, Vitalina Levina, Edwina McGlinn, Chen Davidovich

## Abstract

The Polycomb repressive complex 2 (PRC2) is essential for normal development by maintaining developmental gene repression. PRC2 deposits the repressive chromatin mark H3 lysine 27 tri-methyl (H3K27me3) through a read-write loop that involves direct interactions between PRC2 and H3K27me3. According to current models, the PRC2-H3K27me3 read-write loop is initiated by the PRC2 subunits JARID2 and PALI1 that mimic H3K27me3. However, it is unknown what restricts the PRC2-H3K27me3 read-write loop from spreading H3K27me3 indefinitely. To answer this question, we generated mutant mice where PRC2 subunits cannot mimic H3K27me3. Unexpectedly, the mutations led to delayed Hox genes activation and a homeotic transformation characteristic of a Polycomb gain-of-function in vivo and the spread of H3K27me3 beyond Polycomb domains in stem cells. Collectively, we show that H3K27me3 mimicry evolved to compete against the PRC2-H3K27me3 read-write loop in a process that restrains PRC2 and restricts the spread of Polycomb domains.

**Highlights:**

- H3K27me3 mimicry antagonises Polycomb function in vivo.
- JARID2 and PALI1 synergise to allosterically regulate PRC2 during development.
- H3K27me3 mimicry by JARID2 and PALI1 antagonises PRC2 in stem cells.
- JARID2 and PALI1 mimic H3K27me3 to restrict the spread of Polycomb domains.

## Introduction

When cell type specific genes are repressed, they are packed into facultative heterochromatin, comprised of Polycomb domains. The repressed state of Polycomb domains is maintained by Polycomb repressive complexes ^1^. Polycomb domains are defined by the repressive H3 lysine 27 tri-methyl (H3K27me3) chromatin mark, which is deposited exclusively by the histone methyltransferase Polycomb repressive complex 2 (PRC2) ^2,3^. By maintaining the repressed state of cell-type-specific genes, both PRC2 ^4–6^ and its repressive H3K27me3 mark ^7^ are essential for normal development ^1,8^.

Broad repressive chromatin marks, such as H3K27me3, H3K9me3 and H2AK119ub, are commonly deposited by a “read-write” mechanism: a process where a chromatin modifier binds to its own product (reviewed in ^9,10^). In the case of PRC2, this read-write mechanism ^11^ is especially potent, as it is combined with an allosteric activation ^12^. Specifically, PRC2 is an allosteric enzyme that is largely inactive in its basal state. PRC2 becomes catalytically active when an allosteric effector binds to the allosteric regulatory centre in its core subunit EED ^12^. The best studied allosteric effector of PRC2 is its own product — the H3K27me3 mark ^12^. Upon the deposition of the H3K27me3 mark to chromatin, H3K27me3-modified histone tails can directly bind to the PRC2 subunit EED ^11^ to trigger an allosteric activation of PRC2 ^12^. This mechanism leads to a positive feedback loop that is proposed to maintain the H3K27me3 mark at Polycomb domains when cells are proliferating ^11,12^. Accordingly, pharmacological targeting of the H3K27me3 binding pocket of EED ^13^ or its disruption by mutagenesis ^14^ leads to a loss of H3K27me3 in cells. These perturbation experiments were important to link the allosteric regulatory activity of PRC2 to its function in H3K27me3 deposition in cells. Yet, these experiments perturb the allosteric regulatory activity of PRC2 entirely and, therefore, cannot separate H3K27me3 from the other allosteric effectors of PRC2.

Indeed, three non-histone allosteric effectors of PRC2 were identified, including JARID2 ^15^ and the two paralogous PALI proteins—PALI1 and PALI2 ^16^. JARID2 and PALI1 are accessory subunits of the two distinct types of holo-PRC2 complexes: PRC2.2 and PRC2.1, respectively ^17^. PALI2 was proposed to be an accessory subunit based on sequence homology to PALI1 ^18^. Mechanistically, each of the non-histone allosteric regulators of PRC2 has an effector lysine residue that mimics H3 lysine 27 ^15,16^. These effector lysines—JARID2 K116, PALI1 K1241 and PALI2 K1558—are di- and tri-methylated by PRC2. Once di- and tri-methylated, the effector lysines bind to the allosteric center in EED to switch PRC2 into its stimulated state ^15,16^. Methyltransferase assays using recombinant PRC2-JARID2 complexes demonstrated that JARID2 K116R reduces the activity of PRC2 *in vitro* and JARID2 K116me3 increases it ^19^. Similarly, a mutant PRC2-PALI1 K1241A complex exhibits substantially reduced histone methyltransferase activity *in vitro* ^16^. H3K27me3 mimicry is presumed to be of significance, as it has emerged independently at least twice throughout evolution: H3K27me3 mimicry by JARID2 K116me3 is conserved from fly to human while H3K27me3 mimicry by PALI1 K1241me3 and PALI2 K1558me3 emerged independently in vertebrate species through convergent evolution.

How do the allosteric effectors of PRC2 work together? Early works concluded contradictory roles for JARID2 as either an inhibitory ^20,21^ or stimulatory ^22^ accessory subunit of PRC2. However, later biochemical works found that JARID2 enhances the chromatin binding activity of PRC2 ^23^ and allosterically activates catalysis ^15^. Accordingly, the prevalent paradigm is that the allosteric effector subunits of PRC2, JARID2 and PALI1/2, are methylated by PRC2 to mimic H3K27me3 and then ”jump start” PRC2 during *de novo* deposition of H3K27me3 (reviewed in ^24–33^). Once H3K27me3 is initiated on chromatin, it allosterically activates PRC2 in a self-reinforcing positive feedback loop that maintains H3K27me3 at Polycomb domains. Yet, this model leaves open an unresolved conundrum: what prevents the PRC2-H3K27me3 positive feedback loop from spreading H3K27me3 indefinitely, far beyond Polycomb domains? Moreover, while regulation of PRC2 by the H3K27me3-mimicry activity of PALI1 and JARID2 has been supported based on enzymatic assays *in vitro* and knockout with rescue experiments in cells, this model was never tested *in vivo*. Hence, it is not clear what is the biological relevance of previous models for the regulation of PRC2 by its non-histone effectors through H3K27me3 mimicry.

We set out to determine the physiological and biological relevance of allosteric regulation by the H3K27me3-mimicking subunits of PRC2. To this end, we knocked in allosteric-defective mutations into the endogenous loci of all known PRC2 non-histone effector genes in mouse animals, namely *Jarid2*, *Pali1* and *Pali2*. Herein, we show that the allosteric effector activities of the PRC2 subunits JARID2 and PALI1, but not PALI2, synergise to support normal embryonic development through H3K27me3 mimicry. Yet, contrary to the prevalent paradigm ^24–33^, allosteric defective mouse mutants exhibited skeletal homeotic transformation characteristic of a Polycomb gain-of-function, indicating that H3K27me3 mimicry antagonises Polycomb function. Accordingly, the simultaneous knock-in of allosteric-defective mutations into the endogenous *Jarid2* and *Pali1* loci led to a global gain of H3K27me3 and the spread of Polycomb domains into intergenic regions in mouse embryonic stem cells (ESC). We find that the allosteric regulatory activity of JARID2 and PALI1 is required to restrict H3K27me3 and prevent the spread of Polycomb domains. We propose that H3K27me3 mimicry has repeatedly emerged through evolution to control the spread of Polycomb domains through competition against the PRC2-H3K27me3 read-write loop.

## Results

### The allosteric effector activity of JARID2, but not that of PALI1 and PALI2, is required for mouse development

We aimed to identify the functional consequences of the allosteric regulation of PRC2 by its non-histone effector proteins. To do this, we set out to remove the allosteric regulatory activity of its allosteric effector subunits while keeping these proteins otherwise functional (Figure 1A). Such separation of function (SOF) mutations were identified in JARID2 (K116R) and PALI1 (PALI1 K1241R) within previous biochemical studies ^15,16^. We extended the mutagenesis here also to PALI2 (PALI2 K1558R) based on sequence homology (Figure 1B). These separation of function mutations are minimal lysine-to-arginine perturbations, which only prevent the methylation of the effector lysine and its interactions with EED to prevent the allosteric activation of PRC2 ^15,16^. We generated three separate knock-in mouse lines, in each of them a single separation of function mutation was introduced into one endogenous locus of a non-histone effector, either *Jarid2*, *Pali1* or *Pali2* (Figure 1C). Importantly, unlike a previously studied allosteric mutation in EED, which disrupts the allosteric activity of PRC2 entirely ^14^, in these mouse lines PRC2 can still be subjected to allosteric activation by H3K27me3 (Figure 1A, bottom).

**Figure 1:**
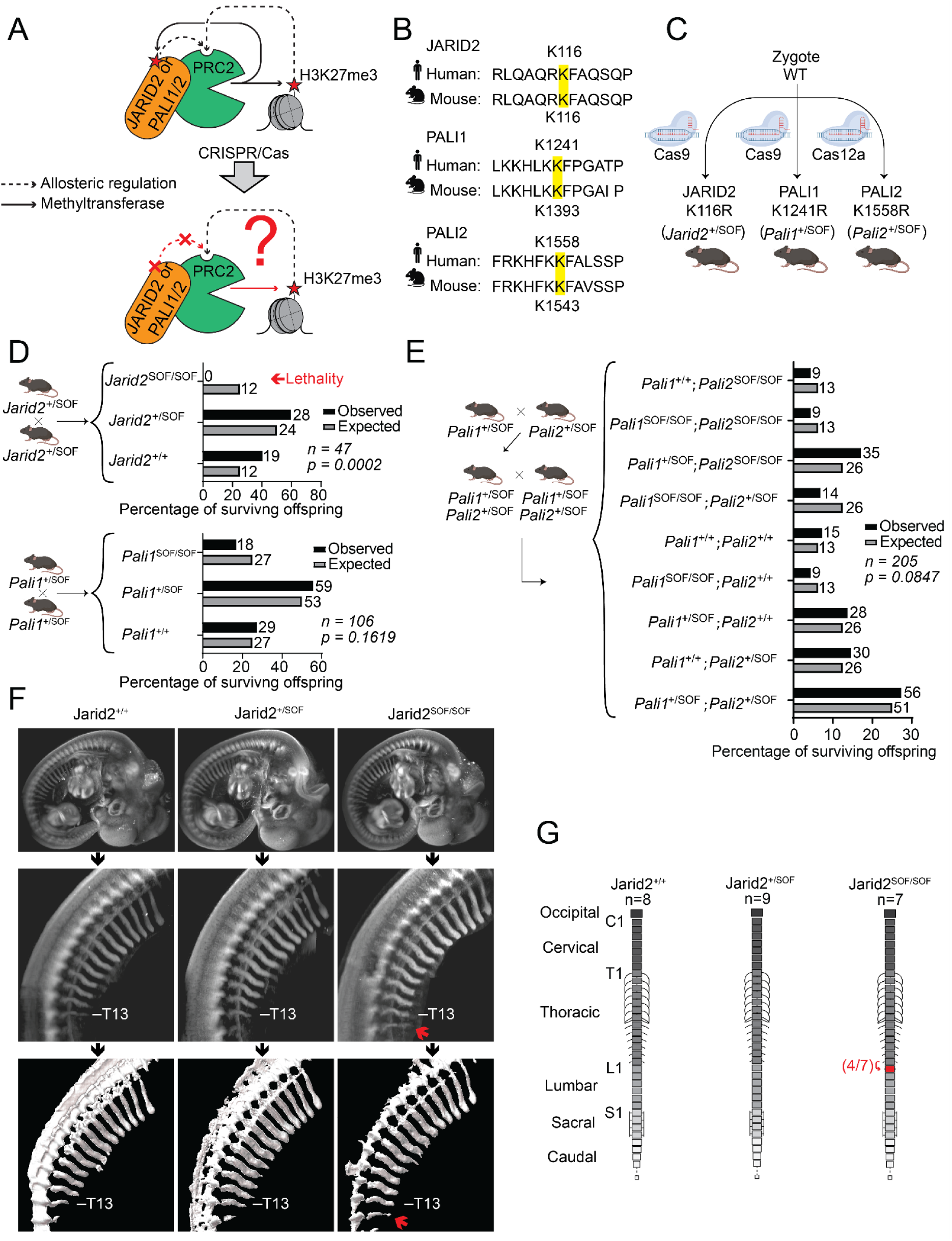
Allosteric defective JARID2, but not PALI1 and PALI2, cause Polycomb gain of function *in vivo*. A. Schematic diagram of allosteric regulation of PRC2. Top: PRC2 methylates an effector lysine—JARID2 K116, PALI1 K1241, PALI2 K1558, or H3K27—that subsequently binds to the allosteric regulatory centre in the PRC2 core to trigger an allosteric activation of methyltransferase. Bottom: Mutating effector lysines in the accessory subunits of PRC2 prevents allosteric activation *in vitro* (dashed red arrow) ^15,16^, but the biological consequences *in vivo* are unknown. B. Alignment of portions of the human and mouse JARID2, PALI1 and PALI2 protein sequences, where the effector lysines are highlighted in yellow. C. Three genetically modified mouse animal lines were generated using CRISPR/Cas knock in of separation-of-function (SOF) mutations into the endogenous *Jarid2*, *Pali1* and *Pali2* genes, as indicated. Each of these mutations include a single lysine-to-arginine mutation of the effector lysine (highlighted in yellow in (B)). D. The percentage of observed (black) and expected (grey) pups that survived to 21 days postnatal as the result of a *Jarid2*^+/SOF^ X *Jarid2*^+/SOF^ crossing. The number of pups from each genotype is indicated. The expected number of pups (adjacent to the grey bars) is rounded to the nearest integer. The total number of pups is indicated by n. The p-value was calculated using a chi-squared test. E. *Pali1^+/SOF^* and *Pali2^+/SOF^* mouse lines were crossed to yield the *Pali1^+/SOF^;Pali2*^+/SOF^ line. Subsequently, *Pali1^+/SOF^;Pali2^+/SOF^* X *Pali1^+/SOF^;Pali2*^+/SOF^ crossing took place and the observed and expected percentage of pups are indicated, as in (D). F. Light sheet fluorescence microscopy of E12.5 embryos immunostained for SOX9. For each genotype, the following are shown: 3D reconstruction of a whole embryo, lateral view (first row), and a separate scan of the same embryo limited to only the region surrounding the thoracic vertebrae, raw data (second row) and 3D rendering (third row). In the second and third rows, only the ribs on one side of the embryo are shown. A second independent comparison is shown in Figure S1A. G. Analysis of homeotic transformation in *JARID2 K116R* mutant embryos. The frequency of homeotic transformations shown in (F) was determined in E14.5 embryos stained with alcian blue. The number of embryos that were examined from each genotype are indicated with n. A schematic diagram representing the mouse vertebral column is shown, where the transformed vertebra is coloured red, with an arrow indicating the direction of the transformation. The number of embryos in which the transformation was observed is indicated beside the corresponding vertebra (in red, left number within the parentheses) and the total number of embryos from the same genotype where that vertebra was visible (in red, right number within the parentheses).

Mice heterozygous for the JARID2 allosteric defective SOF mutation, JARID2 K116R (*Jarid2*^+/SOF^), were viable. Yet, crossing the *Jarid2*^+/SOF^ mice did not produce homozygous mutants (*Jarid2*^SOF/SOF^) that survived to weaning (Figure 1D). JARID2 is essential for mouse development ^34,35^, and our data indicate that its allosteric effector activity is required for this process. While PALI1 is also essential for mouse development ^18^, the allosteric defective PALI1 K1241R homozygous SOF mutant (*Pali1*^SOF/SOF^) animals were viable. Similarly, PALI2 K1558R homozygous SOF mutant mice and PALI1 and PALI2 double homozygous SOF mutant (*Pali1^SOF/SOF^*;*Pali2^SOF/SOF^*) mice were also viable (Figure 1E). These results collectively indicate that allosteric regulation by PALI1 and PALI2 is dispensable during development if *Jarid2* is not perturbed (more below). However, the allosteric regulation of PRC2 by its non-histone effector JARID2 is required for mouse development.

### Allosteric-defective JARID2 leads to a Polycomb gain-of-function phenotype *in vivo*

Past studies using knockout or loss-of-function mutations in various Polycomb group proteins indicate that a hallmark Polycomb loss-of-function phenotype is anterior-to-posterior homeotic transformation (reviewed in ^8^). Therefore, we next assayed for homeotic transformations in the allosteric defective *Jarid2* mouse line. After crossing the *Jarid2^+/SOF^* mouse line, we carried out skeletal imaging on E12.5 and E14.5 embryos using immunofluorescence light sheet microscopy (Figure 1F and Figure S1A) and using alcian blue staining (Figure S1B), respectively. E14.5 was selected, as we find that at this stage most of the embryos of all of the resulting genotypes are still alive. Unexpectedly, we found that approximately half of the *Jarid2^SOF/SOF^* embryos underwent transformation of the first lumbar vertebral element (L1) to a thoracic identity (Figure 1F,G, Figure S1A-C, Supplementary Movies 1-3 and Supplementary Data Table 1). This phenotype was evidenced by the presence of a well formed rib on L1—a posterior-to-anterior transformation—which is characteristic of Polycomb gain-of-function ^8^. Accordingly, Hox gene expression was delayed in the Jarid2^SOF/SOF^ embryos (Figure 2 and Supplementary Data Table 2). Among these, the largest delay was observed for the *Hox10* and *Hox11* genes (Figure 2C, top), which are undergoing activation at this developmental stage (Figure 2C, bottom).

**Figure 2:**
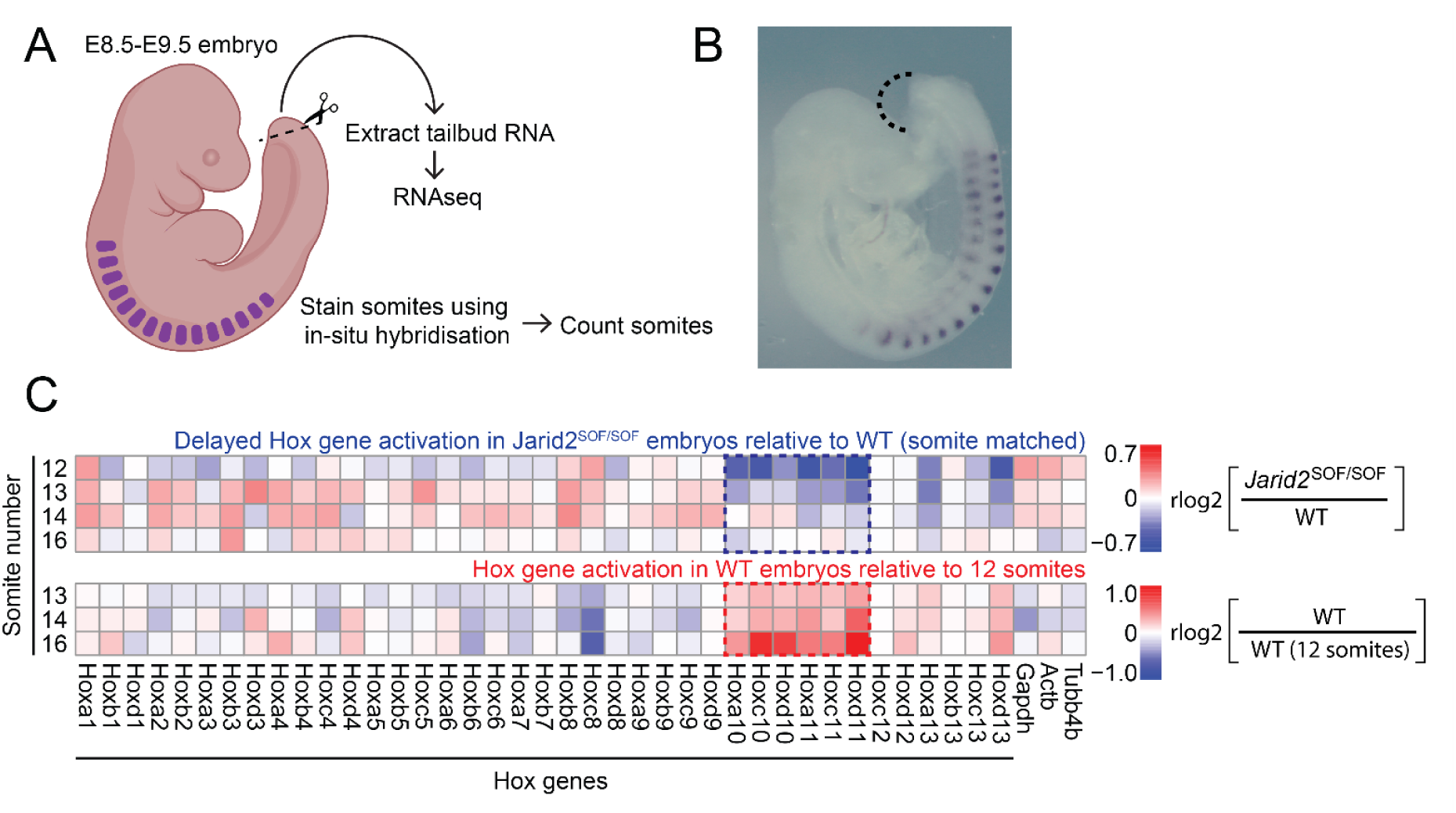
The H3K27me3 mimicry-defective JARID2 mutant leads to delayed transcriptional activation of Hox genes in mouse embryos. A. Schematic of the experiment. Mouse embryos were collected between E8.5 and E9.5, from which RNA was extracted from the tail buds and subjected to RNA-seq. Somites in the body of the embryo were stained using *in situ* hybridisation, then counted to assess the precise developmental stage. B. Representative image of an embryo after in situ hybridisation. The dashed line marks the region from which the tail bud was extracted. C. Top: Heat map showing differential expression of all mouse Hox genes and selected housekeeping genes detected using RNA-seq. Values represent mean rlog2(*Jarid2^SOF/SOF^*) - mean rlog2(WT), calculated using somite-matched embryos from each somite stage as indicated (left). Bottom: Heat map showing Hox gene activation in wild-type embryos, represented as mean rlog2 fold-change relative to the earliest recorded somite stage (12 somites), calculated as mean rlog2(WT) - mean rlog2(WT 12 somites), where mean rlog2(WT) was determined for each of the somite stages 13, 14, or 16, as indicated (left).

We did not observe any consistent homeotic transformation phenotype in any combination of *Pali1* and *Pali2* mutants (Figure S2 and Figure S3C), in agreement with what was previously reported for *Pali1* knockout mice ^18^. These data indicate that the allosteric regulatory activity of JARID2 is essential for mouse development but implies that this activity antagonises Polycomb function.

### The allosteric regulatory activities of PALI1 and JARID2 synergise to sustain embryogenesis

We next wanted to test for genetic interactions between *Jarid2* and *Pali1* or *Pali2*. To test this, we crossed mice heterozygous for both the *Jarid2* and either *Pali1* or *Pali2* SOF mutations (Figure 3A,B). As expected based on our analysis of the single mutants (Figure 1D), there were no surviving mice homozygous for the *Jarid2* SOF allele (*Jarid2^SOF/SOF^*; Figure 3A). Importantly, synthetic lethality occurred when the heterozygous *Jarid2* SOF mutation was present in the background of the homozygous *Pali1* SOF mutation (genotype *Jarid2^+/SOF^*;*Pali1^SOF/SOF^*). Accordingly, only one mouse of the *Jarid2^+/SOF^*;*Pali1^SOF/SOF^* genotype survived to weaning, compared to the expected 20 (Figure 3A, marked with a red arrow). This type of genetic interaction did not occur between *Jarid2* and *Pali2* (Figure 3B).

**Figure 3:**
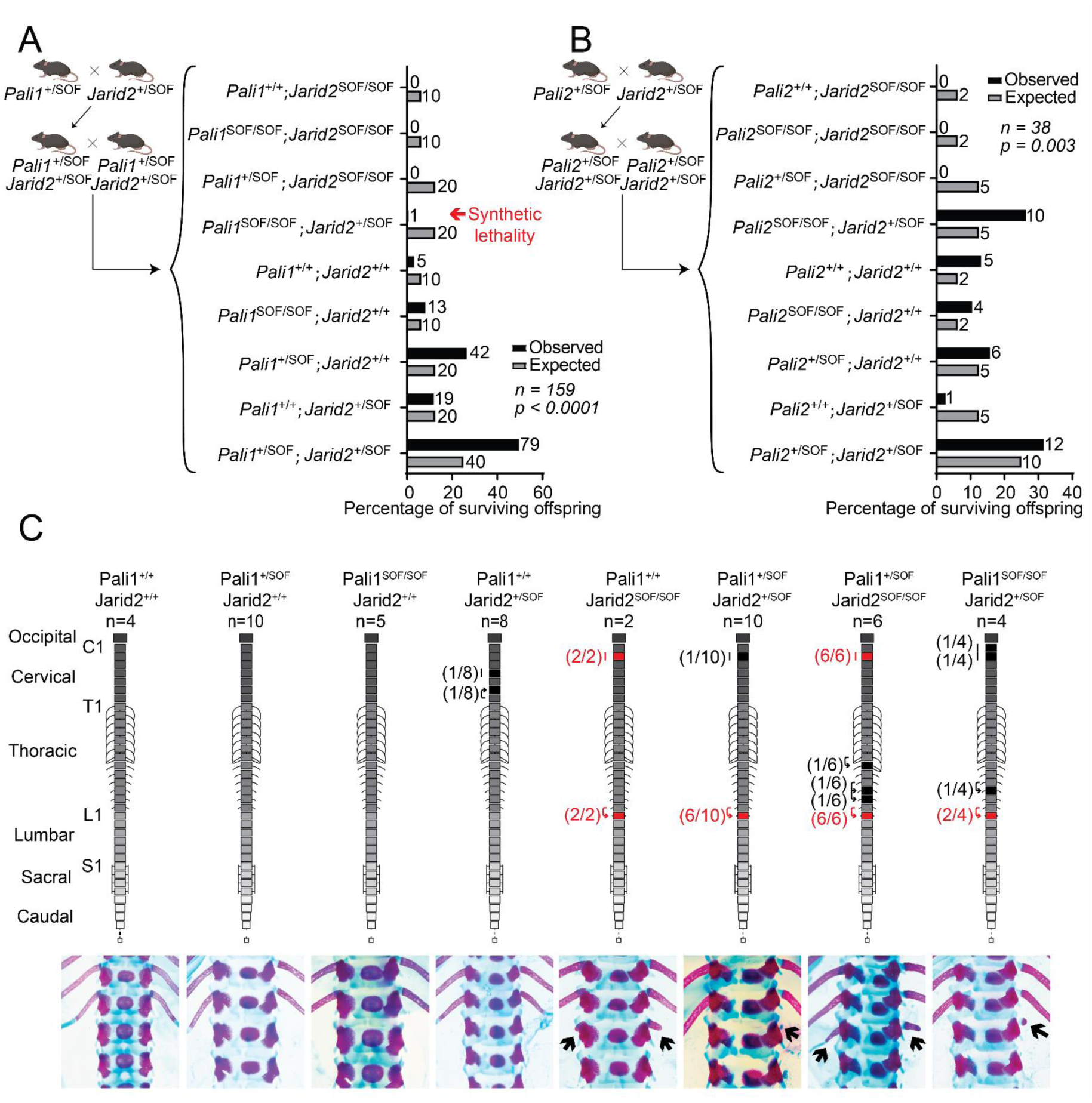
Synergy between allosteric defective JARID2 and PALI1 during mouse embryogenesis. A. Double mutant mice were created by crossing between PALI1 and JARID2 single mutant mice, as indicated. The resulting PALI1 and JARID2 double-heterozygous mutant animals were then crossed with animals of the same genotype. The bar plot represents the percentages of observed (black) and expected (grey) mice of each genotype that survived to 21 days postnatal resulting from these crosses. For each genotype, the observed and expected number of pups are indicated to the right of the corresponding bar (expected number of pups are rounded to the nearest integer). The total number of pups that survived to day 21 postnatal is indicated by n. The p-value was calculated using a chi-squared test. SOF refers to a separation of function mutant: JARID2 K116R or PALI1 K1241R. B. Same as (A), except that PALI2 and JARID2 double mutants were assayed. C. Homeotic transformations in the vertebral column of E18.5 embryos resulting from the *Pali1^+/SOF^;Jarid2^+/SOF^ x Pali1^+/SOF^;Jarid2^+/SOF^* crossing. Top: A schematic diagram representing the mouse vertebral column, including the occipital bone up to the 4^th^ caudal vertebra (no transformations were observed in the caudal vertebrae). Vertebrae in which homeotic transformation or malformation were observed with a frequency of at least 50% are coloured red, or black if it was observed with a frequency of less than 50%. The direction of homeotic transformations are indicated by an arrow. The number of embryos that the transformation was observed in is given next to the corresponding vertebra. Bottom: Representative photos of the T12-L2 vertebrae showing the L1 to T13 transformation, indicated with arrows.

For genotypes that were identified as lethal prior to weaning, we characterised the axial skeleton at embryonic day E18.5. This analysis revealed consistent posterior-to-anterior homeotic transformations (Figure 3C, Figure S3A and Supplementary Data Table 1) and an increase in the total number of vertebrae (Figure S3B). These posterior-to-anterior homeotic transformations are in agreement with the posterior-to-anterior transformation identified in the *Jarid2^SOF/SOF^* embryos (Figure 1F,G and Figure S1) and are in agreement with a Polycomb gain-of-function phenotype ^36–45^. While some of the transformations appeared only in certain genotypes and with a low penetrance, the most prevalent homeotic transformation is the one we identified in the *Jarid2*^SOF/SOF^ embryos (Figure 1F,G and Figure S1): a posterior-to-anterior transformation of L1 to T13 (Figure 3C). Crucially, this phenotype was also seen in about half of the *Jarid2* heterozygotes when that mutation was present on the background of at least one mutant PALI1 allele (specifically, *Jarid2^+/SOF^*;*Pali1^+/SOF^* or *Jarid2^+/SOF^*;*Pali1^SOF/SOF^* in Figure 3C). This is important, as it indicates a synergy between the allosteric regulatory activities of JARID2 and PALI1 ^45^. In the context of this work, we adhere to the definition of synergy as ‘occurs when the contribution of two mutations to the phenotype of a double mutant exceeds the expectations from the additive effects of the individual mutations’ ^46^. Here, the synergy between JARID2 K116 to PALI1 K1241 has an apparent antagonistic effect on Polycomb function *in vivo*.

### Allosteric-defective mutations in the PRC2 accessory subunits increases H3K27me3 in cells

The unexpected observation that JARID2 K116R and PALI1 K1241R mutations exhibited Polycomb gain of function phenotypes *in vivo* indirectly suggests a PRC2 gain of function. To directly test if these mutations enhance the activity of PRC2 in cells, we knocked-in either the JARID2 K116R or the PALI1 K1241R homozygous mutations into the endogenous *Jarid2* or *Pali1* loci in mouse embryonic stem cell (mESC) lines using CRISPR/Cas9 mediated gene editing (*Jarid2^SOF/SOF^* and *Pali1^SOF/SOF^* cell lines, respectively; in Figure 4A). We then knocked-in the homozygous JARID2 K116R mutation into the endogenous *Jarid2* locus within the *Pali1^SOF/SOF^* cell line, to generate the homozygous double mutant *Jarid2^SOF/SOF^*;*Pali1^SOF/SOF^* (Figure 4A, bottom).

**Figure 4:**
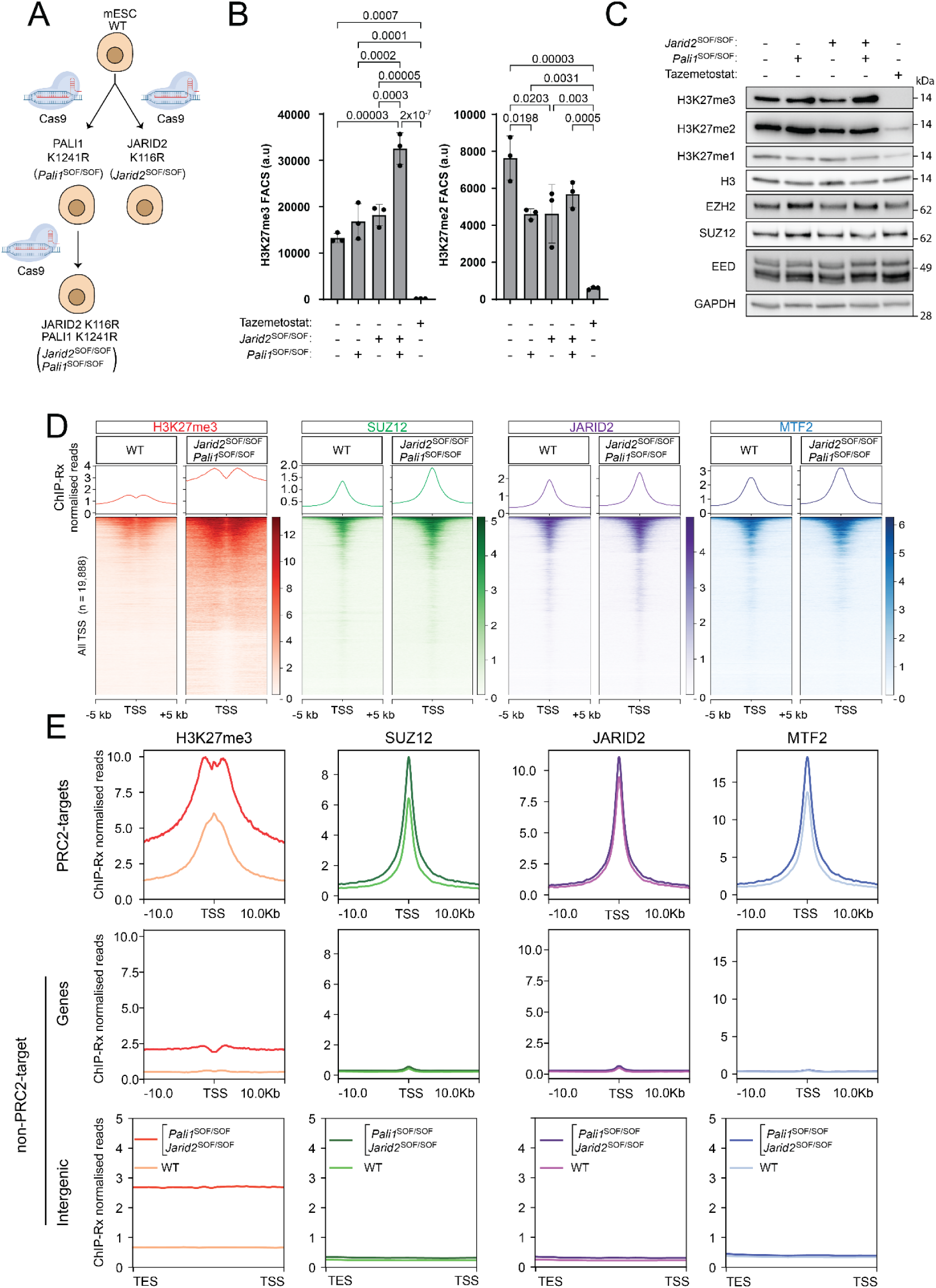
Allosteric defective JARID2 and PALI1 lead to increased H3K27me3 in mESC. A. Two genetically modified mESC lines were generated using CRISPR/Cas knock-in of the indicated separation-of-function (SOF) mutations into the endogenous *Jarid2* and *Pali1* loci. Subsequently, a double mutant cell line was generated by knocking in the JARID2 K116R mutation into the *Jarid2* locus within the PALI1 K1241R cells. All knock-ins were homozygous. B. Fluorescence-activated cell sorting (FACS) on cells possessing mutations or Tazemetostat treatment as specified. The bar charts show the mean of the median fluorescence signal and error bars represent the standard error across three independent biological replicates that were harvested on different days. Statistical significance was calculated using a one-way ANOVA with a post-hoc Tukey’s test to correct for multiple comparisons. Non-significant comparisons are not shown. C. Western blots on whole cell lysate of cells possessing the indicated mutations or Tazemetostat treatment using antibodies as specified. See Figure S5A,B for two more replicates and a densitometry analysis. D. Quantitative ChIP-seq (ChIP-Rx) was carried out using cell lines and antibodies as indicated and heatmaps are centred on all transcription start sites. ChIP-Rx was performed using antibodies for H3K27me3 (red), SUZ12 (green), JARID2 (purple), and MTF2 (blue). E. Average ChIP-Rx signal profiles at all transcription start sites at PRC2-target genes, non-PRC2-target genes, or intergenic regions as indicated. Colour keys are presented in the bottom plots. For a second independent ChIP-Rx replicate that was carried out on a different day see Figure S4A,B.

The *Jarid2^SOF/SOF^*;*Pali1^SOF/SOF^* double mutant cell line exhibited an approximately two-fold increase in H3K27me3 globally, as detected by flow cytometry (Figure 4B, Figure S5C) and immunoblotting (Figure 4C, Figure S5A,B). Accordingly, quantitative ChIP-seq with a reference exogenous genome ^47^ revealed substantial increase of H3K27me3 at transcription start sites (TSS) in the *Jarid2^SOF/SOF^*;*Pali1^SOF/SOF^* cells with respect to the wild type (Figure 4D and Figure S4). H3K27me3 was also increased in the *Jarid2^SOF/SOF^* cells, but not to the same level as *Jarid2^SOF/SOF^*;*Pali1^SOF/SOF^* (Figure S6A,B). Increased H3K27me3 was observed in PRC2-target genes, defined by their association with SUZ12 peaks in the wild-type cells, but also at non-PRC2 target genes and intergenic regions (Figure 4E and Figure S4). The gain in H3K27me3 occurs at a similar level in all H3K27me3-marked genes, regardless of how broad H3K27me3 peaks were in the wild-type cells (Figure S6C). We also did not record changes to H3K4me3 methylation in the *Jarid2^SOF/SOF^*;*Pali1^SOF/SOF^* cells (Figure S6D), suggesting that bivalent chromatin remained the same. These observations imply that even in the presence of H3K27me3-mimicry defects, PRC2 still obeys the same rules as in normal cells, with the exception that it is now more active in canonical Polycomb domains.

It is possible that the mutations JARID2 K116R and PALI1 K1241R enhance the activity of PRC2 by ensuring that the allosteric regulatory site is always available to bind H3K27me3. This is consistent with *in vitro* HMTase assays, indicating that a H3K27me3 histone tail peptide appears to be a better allosteric effector than JARID2 K116me3 and PALI1 K1241me3 (Supplementary Figure S6E). Accordingly, the mutations did not substantially change nuclear localisation or the apparent stability of PALI1 and JARID2 (Figure S4E-F), pointing to changes in the intrinsic activity of PRC2 as a determinant.

The increase in H3K27me3 was accompanied by a smaller but reproducible increase of the core PRC2 subunit SUZ12, the PRC2.2 subunit JARID2 and the PRC2.1 subunit MTF2 (Figure 4E and Figure S4). This mirrors observations in previous reports, where histone methyltransferase-defective PRC2 mutants exhibited reduced chromatin occupancy in cells ^48–50^. This phenotype is different from the JARID2 knockout phenotypes, which did not lead to change in global H3K27me3 ^51^ and on average slightly reduced H3K27me3 at PRC2 target genes in mECS, either undifferentiated ^52^ or differentiated ^49^. We also demonstrated that such a large increase of global H3K27me3 is unlikely to occur merely because of the clonal selection (Figure S7B-D). These observations indicate that the same JARID2 and PALI1 allosteric defective mutants that lead to a Polycomb gain-of-function phenotype *in vivo* (Figure 1-3) also increase the histone methyltransferase activity of PRC2 in cells (Figure 4).

The mutations did not change the number or size of RING1B foci, sometimes referred to as Polycomb bodies (Figure S8A-D). Specifically, stimulated emission depletion (STED) super resolution microscopy indicated that the size of RING1B foci remained around 1×10^-2^ µm^2^, which is close to the detection limit of the imaging approach (∼2.8×10^-3^ µm^2^ with our imaging parameters). ChIP-seq with a reference exogenous genome did not detect a substantial change to RING1B on chromatin in the *Jarid2^SOF/SOF^*;*Pali1^SOF/SOF^* cells (Figure S8E). However, H2AK119ub is reduced in the mutant cells (Figure S8F). While future studies will likely determine the reason for the reduced H2AK119ub deposition, a contributing factor could be that the same number of PRC1 molecules have to deposit H2AK119ub to larger Polycomb domains in the *Jarid2^SOF/SOF^*;*Pali1^SOF/SOF^* cells.

### The allosteric regulatory activity of PRC2 accessory subunits restricts the spread of Polycomb domains

In order to gain mechanistic insights on how the allosteric regulatory subunits of PRC2 reduces its histone methyltransferase activity, we analysed the spread of H3K27me3 (Figure 5A-D). We found that in the wild-type cells H3K27me3 is mostly restricted to CpG islands, but in the *Jarid2^SOF/SOF^*;*Pali1^SOF/SOF^* cells H3K27me3 has spread tens of kilobase pairs further away from the nucleation sites in both directions (Figure 5A). SUZ12, MTF2 and JARID2 are still localised to the same sites on chromatin in both the wild-type and mutant cell lines (Figure 5A; antibody for the PRC2.1 subunit MTF2 was used as there is no suitable ChIP antibody for PALI1). This indicates that in the mutant cells, H3K27me3 is spread away from Polycomb domains in a process that does not involve a spread of PRC2 beyond its natural nucleation sites. In the *Jarid2^SOF/SOF^*;*Pali1^SOF/SOF^* cells, H3K27me3 was increased both inside and outside of PRC2-target TSS (Figure 4E). Yet, differential analysis of the ChIP-Rx coverage indicated that the gain of H3K27me3 was comparatively higher around PRC2 binding sites than inside them (Figure 5B-C). Accordingly, substantially less ChIP-Rx H3K27me3 reads from within the mutant cell lines, in comparison to the wild type, were mapped to SUZ12 peaks (Figure S4C). Conversely, differential analysis of SUZ12 ChIP-Rx indicated an increase at the PRC2 target TSS, defined using the wild-type cells, but not beyond TSS (Figure 5C, in green, and Figure 5D). Collectively, these data indicate that JARID2 and PALI1 synergise to restrain PRC2 *in vivo* by restricting the spread of H3K27me3 beyond Polycomb domains.

**Figure 5:**
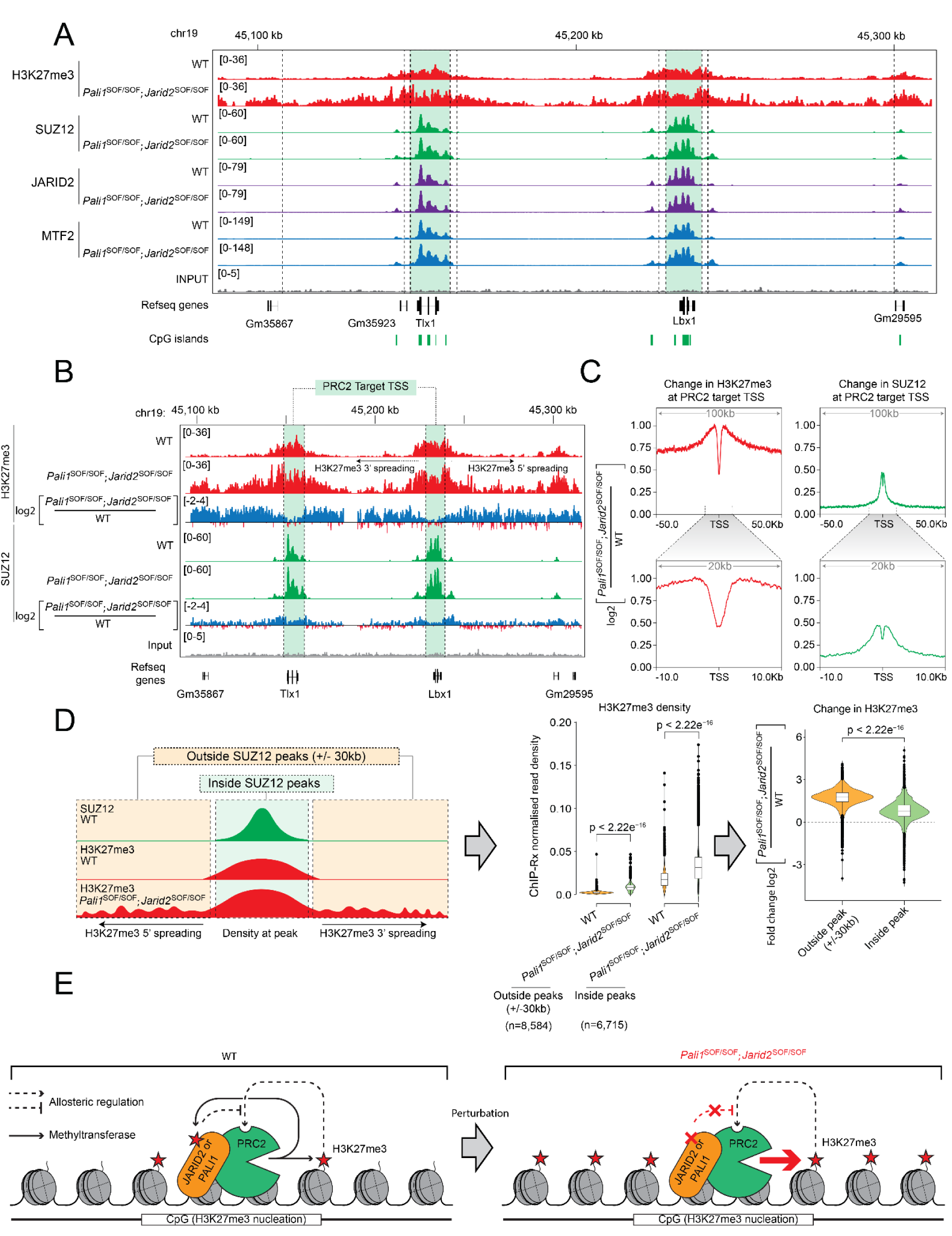
The allosteric regulatory activity of JARID2 and PALI1 is required to restrict the uncontrolled spread of Polycomb domains. A. Gene track over a representative locus showing ChIP-Rx signals from cell lines and antibodies as indicated. PRC2 nucleation sites are defined by the SUZ12 peaks in the wild-type cells and are highlighted in green. B. Gene tracks comparing H3K27me3 (top) or SUZ12 (bottom) in the PALI1 K1241R and JARID2 K116R double mutant cells with respect to the wild type. Shown from top to bottom for each antibody: quantitative ChIP-seq (ChIP-Rx) signal in wild-type cells, ChIP-Rx signal in the double mutant cells, wild type ChIP-Rx divided by the double mutant ChIP-Rx. In the latter case, higher or lower ratios are coloured blue or red, respectively. C. The average change in H3K27me3 or SUZ12 at all PRC2 target sites. Data is plotted as a log2 ratio of reads in the PALI1 K1241R and JARID2 K116R double mutant to wild type. Antibodies are indicated. Different ranges around TSS are presented in the top and bottom plots, as indicated. D. Left: A schematic diagram showing the change in H3K27me3 localisation in the mutant with respect to the wild-type cells. Middle: The H3K27me3 ChIP-Rx read density inside and within 30 kbp outside of SUZ12 peaks (highlighted within the illustration on the left in green and yellow, respectively). The median is shown with a dashed line. Right: The change in the H3K27me3 ChIP-Rx read density inside and outside of peaks in the mutant relative to the wild-type cells. Statistical significance was assessed by a Mann-Whitney U test and p values are indicated. E. Schematic diagram showing a model for allosteric regulation of PRC2 by JARID2 and PALI1. In wild-type cells, PRC2 methylates JARID2 K116 and PALI1 K1241, which then allow them to compete against H3K27me3 for binding to the regulatory center in PRC2. This process breaks the PRC2-H3K27me3 read-write positive feedback loop to retain normal levels of H3K27me3 at CpG islands. In the allosteric defective mutant cell line (right), the mutant PALI1 and JARID2 can no longer compete with H3K27me3. As a result, the read-write loop is no longer controlled, leading to high levels of H3K27me3 at CpG islands and a spread of H3K27me3 beyond Polycomb domains. Across the figure, H3K27me3, SUZ12, JARID2 and MTF2 ChIP-seq data are marked in red, green, purple and blue, respectively.

## Discussion

Allosteric regulation is a central tenet in chromatin biology, activating chromatin modifiers in the presence of the right effector to enable context-specific regulation ^25,53^. However, little is known about how multiple allosteric effectors synergise to regulate the activity of a given chromatin modifier *in vivo*. Previous works revealed that the methylation of effector lysines in JARID2 and PALI1 stimulates PRC2 *in vitro* ^15,16^. Based on knockout with rescue experiments, these works concluded that JARID2 K116me2/3 and PALI1 K1241me2/3 stimulate, or “jump start”, PRC2 to deposit H3K27me3 *de novo* during gene repression ^15,16^. This “jump start” model ^24^ is largely accepted ^24–33^ and possibly correct. Yet, using a set of knock-in mouse lines and stem cells we find that the main biological role of the effector lysines in JARID2 and PALI1 is the opposite to what was assumed so far: they largely restrain PRC2 *in vivo*, not enhancing its activity. Intrinsic lysine methylation has already been recognised as an important mechanism to control positive feedback loops in chromatin modification, demonstrated for H3K9 methylation in yeast ^54^ and more recently in mammalian cells ^55^. Our new data, when contrasted with previous works ^11,12,15,16^, allow piecing together a model that explains how multiple allosteric effectors synergise to regulate PRC2 in cells:

During *de novo* H3K27me3 deposition, JARID2 K116me2/3 and PALI1 K1241me2/3 may allosterically activate PRC2, as previously proposed ^15,16^. Once H3K27me3 has been deposited to Polycomb domains, PRC2 molecules that bind there use their regulatory subunit EED to interact with the pre-deposited H3K27me3 ^11^. The interaction between H3K27me3 and PRC2 triggers the allosteric activation of PRC2 ^12^ and, at the same time, increases the affinity of PRC2 to chromatin ^11^. Now, the allosterically activated PRC2 introduces more H3K27me3 to chromatin. Importantly, this process can take place even if not all the JARID2 and PALI1 molecules in the cell are methylated, as these proteins have a strong chromatin binding activity disregarding their effector lysines ^16,23,49,56^. Since the chromatin-bound PRC2 is now allosterically activated, it can occasionally also methylate JARID2 K116 ^15^ or PALI1 K1241 ^16^. The affinity of the effector lysines JARID2 K116me2/3 and PALI1 K1241me2/3 for EED is about 3- to 5-fold higher than that of the H3K27me3 histone tails to EED ^15,16^. Moreover, JARID2 and PALI1 are already part of the holo-PRC2 complex ^15,16^, which increases the effective concentration of JARID2 K116me2/3 and PALI1 K1241me2/3 around EED. Hence, as soon as an effector lysine in JARID2 or PALI1 becomes di- or tri-methylated, it blocks the allosteric regulatory site in EED and prevents H3K27me3 tails from binding there. JARID2 K116me2/3 and PALI1 K1241me2/3 can still allosterically activate PRC2, although they do not directly contribute to chromatin binding ^15,16^. Conversely, the interactions between H3K27me3 and EED contribute to both allosteric activation ^12^ and chromatin binding ^11^, because H3K27me3 is part of the chromatin. This likely makes H3K27me3 a stronger activator of PRC2 compared to the effector lysines in PALI1 and JARID2.

This effect might be further enhanced if the H3K27 tri-methyl lysine is a better allosteric effector than its mimicking lysins within JARID2 and PALI1, which is the case *in vitro* (Supplementary Figure 6E). This model fits with the antagonistic activity of the JARID2 and PALI1 effector lysines in cells (Figure 4-5) and *in vivo* (Figure 1-3) and also with their stimulatory activity *in vitro* (Figure S6E and ^15,16^).

We propose that JARID2, PALI1 and H3K27me3 work together to maintain a healthy level of H3K27me3 on chromatin. A high level of H3K27me3 would increase the instances where PRC2 is allosterically activated. Sustained allosteric activation of PRC2 would eventually lead to the di/tri-methylation of JARID2 and PALI1 and the subsequent interference with the PRC2-H3K27me3 positive feedback loop. This mechanism is unlikely to take place in active genes, where the H3K4me3 and H3K36me3 histone mark inhibit PRC2 ^57^ and H3K4me3 may compete against JARID2 K116me2/3 for binding to EED there ^58^. When JARID2 K116 and PALI1 K1241 are mutated, the PRC2-H3K27me3 positive feedback loop cannot be restrained, leading to increased H3K27me3 at CpG islands (Figure 4) and eventually to its spread away from Polycomb domains (Figure 5).

Ectopically expressed PALI2 binds to PRC2 in cells ^18^ and a polypeptide that includes PALI2 K1558me3 is sufficient to stimulate PRC2 *in vitro* ^16^. However, our data indicate that PALI2 K1558 is dispensable for mouse development, even on the background of the PALI1 K1241R mutation (Figure 3). At this time, we are not aware of direct evidence for functional interactions between the endogenously expressed PALI2 and PRC2 in cells.

Intriguingly, mutations in and around JARID2 K116 and PALI1 K1241, including recurrent mutations, have been observed in cancer. Examples are JARID2 K116T ^59^ and K116N ^60^ in carcinoma and PALI1 P1243S ^61–63^ and P1243A ^62^ in melanoma ^64,65^. If these cancer-associated mutations interfere with the binding of the JARID2 or PALI1 effector lysines to EED, then they may confer a PRC2 gain-of-function, as with JARID2 K116R and PALI1 K1241R (Figure 1-5). PRC2 gain-of-function is commonly associated with poor prognosis in cancer ^66^ and in some cases can be targeted by the pharmacological inhibition of PRC2 ^67^.

We show that the allosteric regulatory activity of the accessory subunits of PRC2 is required for embryonic development and that allosteric-defective JARID2 and PALI1 lead to homeotic transformations (Figure 1 and 3), a hallmark Polycomb gain-of-function phenotype. Anterior-to-posterior axial skeletal transformations have been identified as a loss-of-function phenotype in mice for various Polycomb group proteins ^36–45^. To our knowledge, no homeotic transformation has been reported in studies where *Pali1* ^18^ or *Jarid2* ^35,68–70^ were knocked-out in mice. Perinatal or embryonic lethality was observed in the case of both PALI1 ^18^ and JARID2 ^35^ knock-out (reviewed in ^34^), and the latter also led to organ or skeletal development defects ^35,68–70^. However, it was impossible to link these previously-identified phenotypes directly to Polycomb function, especially in the case of PALI1 that also binds to the H3K9me2 methyltransferase G9A ^18^. Moreover, regions of JARID2 and PALI1, external to their effector lysines, enhance the affinity of PRC2 for nucleosomes ^16,23^ and are required for chromatin binding by PRC2 ^49,56^. Therefore, knockout experiments cannot link a specific biochemical function of PALI1 or JARID2 to the observed phenotype. Our results indicate that the allosteric regulatory activities of JARID2 (Figure 1) and PALI1 (Figure 3) are required for normal development and reveal a synergy between them (Figure 3) in antagonising Polycomb function (Figure 3-5).

PALI1 is a vertebrate-specific protein ^18^ while JARID2, including K116, is conserved from fly to human ^15^. Their mechanistic resemblance *in vitro* led to the proposal that PALI1 emerged through a convergent evolution to mimic JARID2 at the molecular level ^16^. Our discovery of genetic interactions between JARID2 K116 and PALI1 K1241 indicate that JARID2 and PALI1 synergise to allosterically regulate PRC2 *in vivo* through H3K27me3 mimicry. We propose that H3K27me3 mimicry by JARID2 evolved to restrict the read-write PRC2-H3K27me3 positive feedback loop of PRC2.2 in metazoans while PALI1 emerged in vertebrate species to carry out this function in the context of PRC2.1. Collectively, JARID2 and PALI1 synergize to buffer the PRC2-H3K27me3 positive feedback loop, allowing PRC2 to sustain H3K27me3 at Polycomb domains without expanding beyond them.

## Limitations of the Study

In this work, we did not probe for changes to 3D genome architecture. Such analysis may reveal how chromatin folding might facilitate the spread of H3K27me3 despite PRC2 remaining tethered to CpG islands. While we detected the chromatin occupancy of RING1B using quantitative ChIP-seq, we did not probe for cPRC1- and vPRC1-specific subunits. Therefore, we cannot exclude the possibility that changes to the chromatin occupancy of specific types of PRC1 complexes may occur in mutant cells defective in H3K27me3 mimicry. We also did not probe for PALI1 directly in our ChIP-seq experiments, given the lack of suitable antibodies. Instead, we used anti-MTF2 antibodies to detect PRC2.1. While our STED super resolution microscopy did not detect changes to the size of RING1B foci in the *Jarid2^SOF/SOF^*;*Pali1^SOF/SOF^* cells, we cannot exclude the possibility that changes to the size of such foci may occur in mECS but are beyond the resolution limit of light microscopy.

## Acknowledgements

We thank the support of the Monash FlowCore, Monash University Palaeontology Lab, and the MASSIVE HPC facility. The mutant mouse animals and cells were produced via CRISPR genome editing by the Monash Genome Modification Platform (MGMP), Monash University, as a node of Phenomics Australia. Phenomics Australia is supported by the Australian Government Department of Education through the National Collaborative Research Infrastructure Strategy, the Super Science Initiative and the Collaborative Research Infrastructure Scheme. S.C.A. was supported through an Australian Government Research Training Program (RTP) Scholarship. C.D. was an EMBL-Australia Group Leader and a Sylvia and Charles Viertel Senior Medical Research Fellow, is an ARC Future Fellow (FT240100821), and acknowledges support from the ARC (DP190103407) and the NHMRC (APP1162921, APP1184637, APP2011767, APP2020900 and APP2046466). This research was funded partially by the Victoria State Government through mRNA Victoria (C.D.).

## Authors contributions

S.C.A. and C.D. conceptualised the project, S.C.A, M.B., Y.K, Y.C, M.M., J.M., Z.L., S.N., and E.M. carried out experiments and investigated, S.C.A., M.B., V.L. and C.D. managed mouse colonies and animal ethics, J.R. generated cell lines and mouse animal lines, Z.L. produced reagents, S.C.A and E.H. carried out bioinformatic analyses, Y.C, J.H-L, and E.M. provided training and input into experimental design, J.H-L created the image analysis pipeline, C.D. and E.M. supervised, S.C.A. and C.D. wrote the original draft and all authors reviewed and edited the manuscript.

## Data and materials availability

Next generation sequencing data (ChIP-Rx and RNA-seq) are available under GEO accession number GSE278724.

---

**Supplementary Figure 1:**
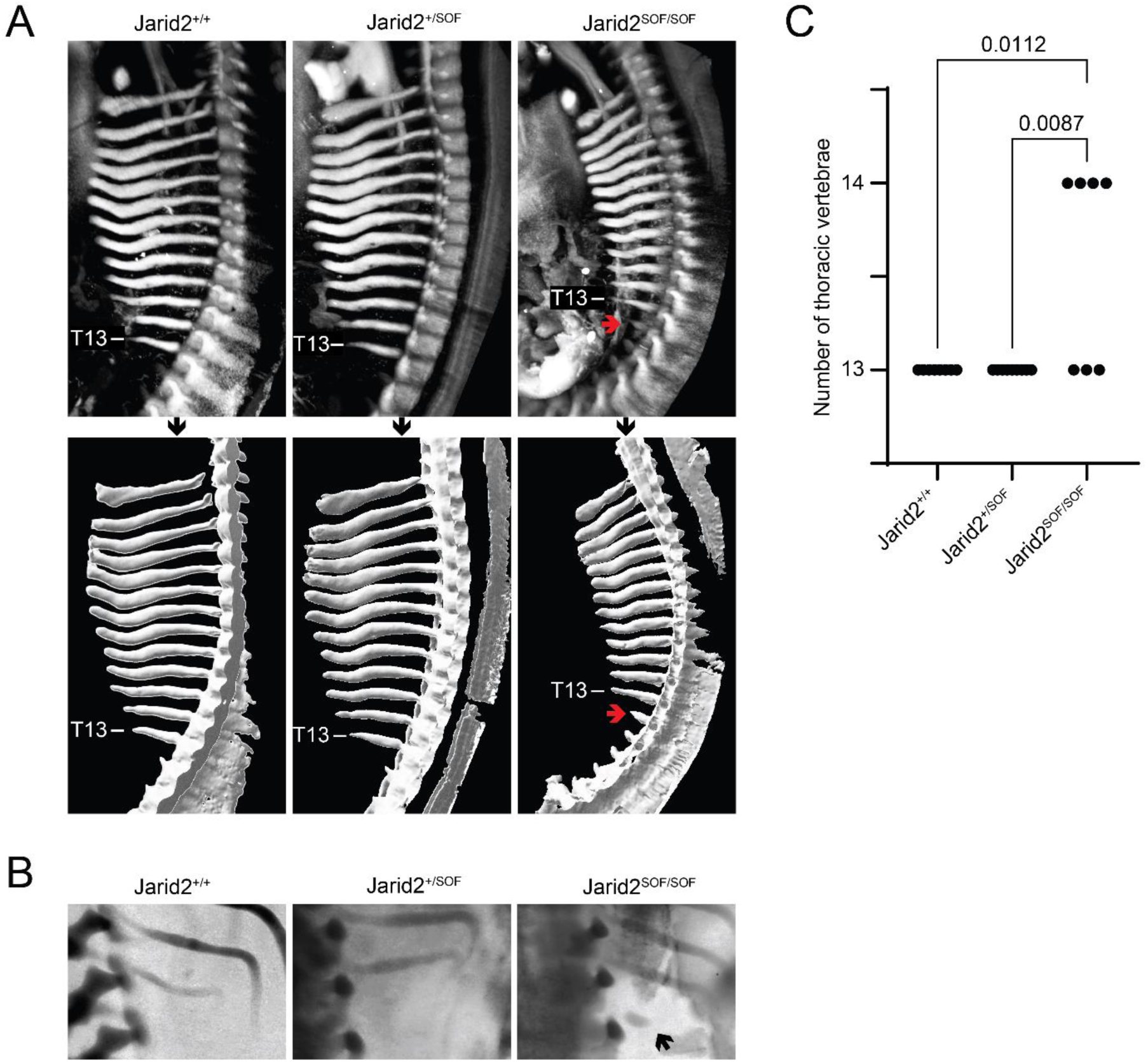
*Jarid2^SOF/SOF^* causes posterior to anterior homeotic transformation in mouse. A. Light sheet fluorescence microscopy of the region surrounding the thoracic vertebrae in E12.5 embryos stained for SOX9. A 3D rendering from surface segmentation is shown in the second row. The homeotic transformation L1 to T13 is indicated with an arrow. Presented is another independent comparison from the same experiment as in Figure 1F. B. Representative images of the E14.5 embryos corresponding to the analysis in Figure 1G. All embryos in this analysis were stained with alcian blue and imaged in a dissecting microscope with white light. Representative embryos from each of the genotypes are shown, where the homeotic transformation L1 to T13 is indicated with an arrow. C. Total number of thoracic vertebrae for the embryos that were part of the experiment, as shown in Figure 1G. Each dot represents the number of thoracic vertebrae in one embryo. Genotypes are indicated. A Kruskal-Wallis test with Dunn’s test for multiple comparisons was used to test for significance between all combinations of genotypes. Only statistically significant p values are presented.

**Supplementary Figure 2:**
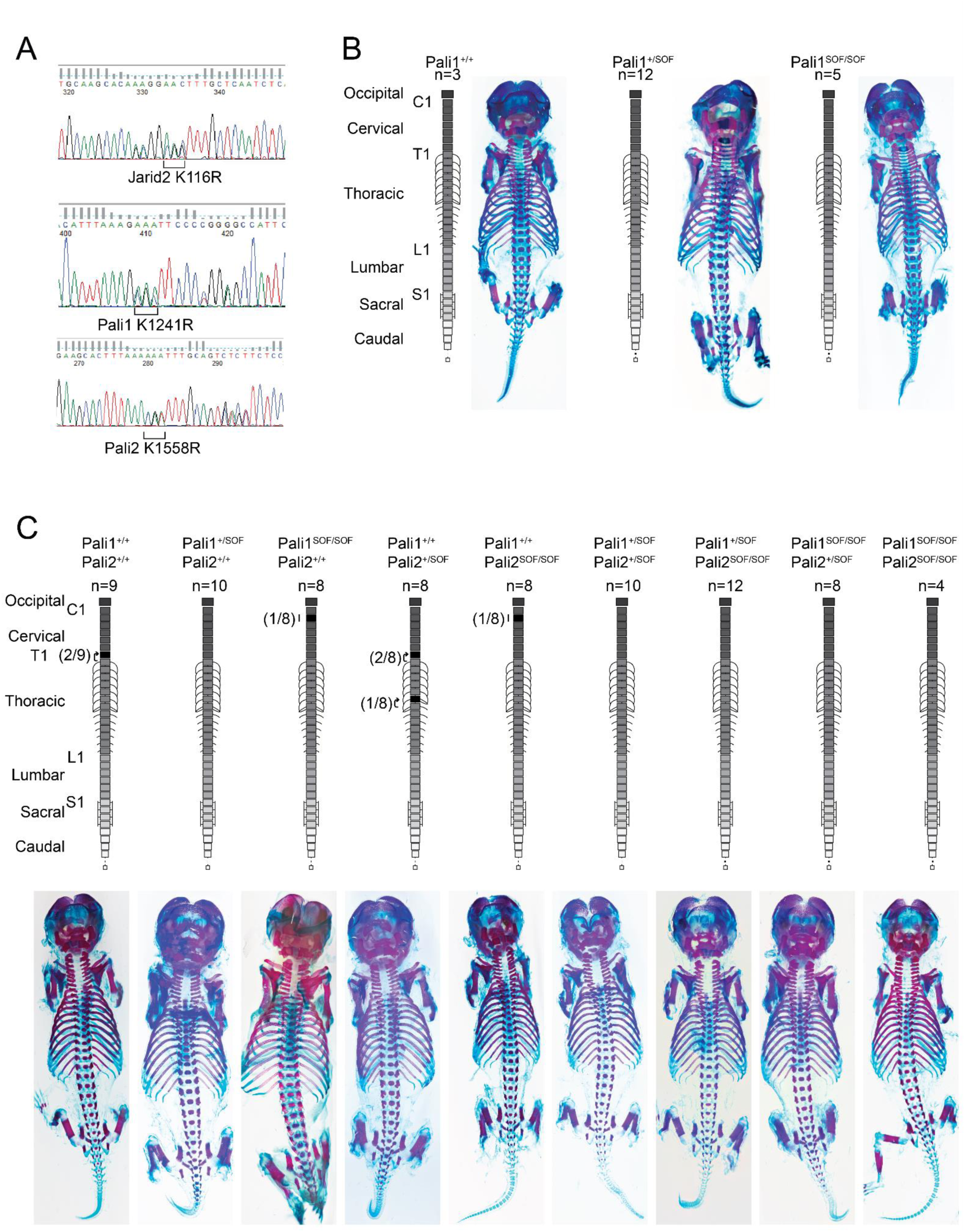
PALI1 K1241 and PALI2 K1558 are dispensable for skeletal patterning. A. Sanger sequencing traces confirming the correct knock-in heterozygous mouse lines that were generated in this study. The codon corresponding to the separation of function mutation is annotated below. B. No homeotic transformation or other defects were observed in embryonic day 18.5 (E18.5) skeletons resulting from *Pali1^+/SOF^* x *Pali1^+/SOF^* crossing. The number of embryos that were examined from each genotype is indicated by n. A representative photo of each genotype is shown. C. The same as (B), for *Pali1^+/SOF^;Pali2^+/SOF^ x Pali1^+/SOF^;Pali2^+/SOF^* crossings. Vertebrae in which homeotic transformation or malformation were observed are coloured black, with the direction of homeotic transformations indicated by an arrow. The number of embryos that the transformation was observed in is given next to the corresponding vertebra.

**Supplementary Figure 3:**
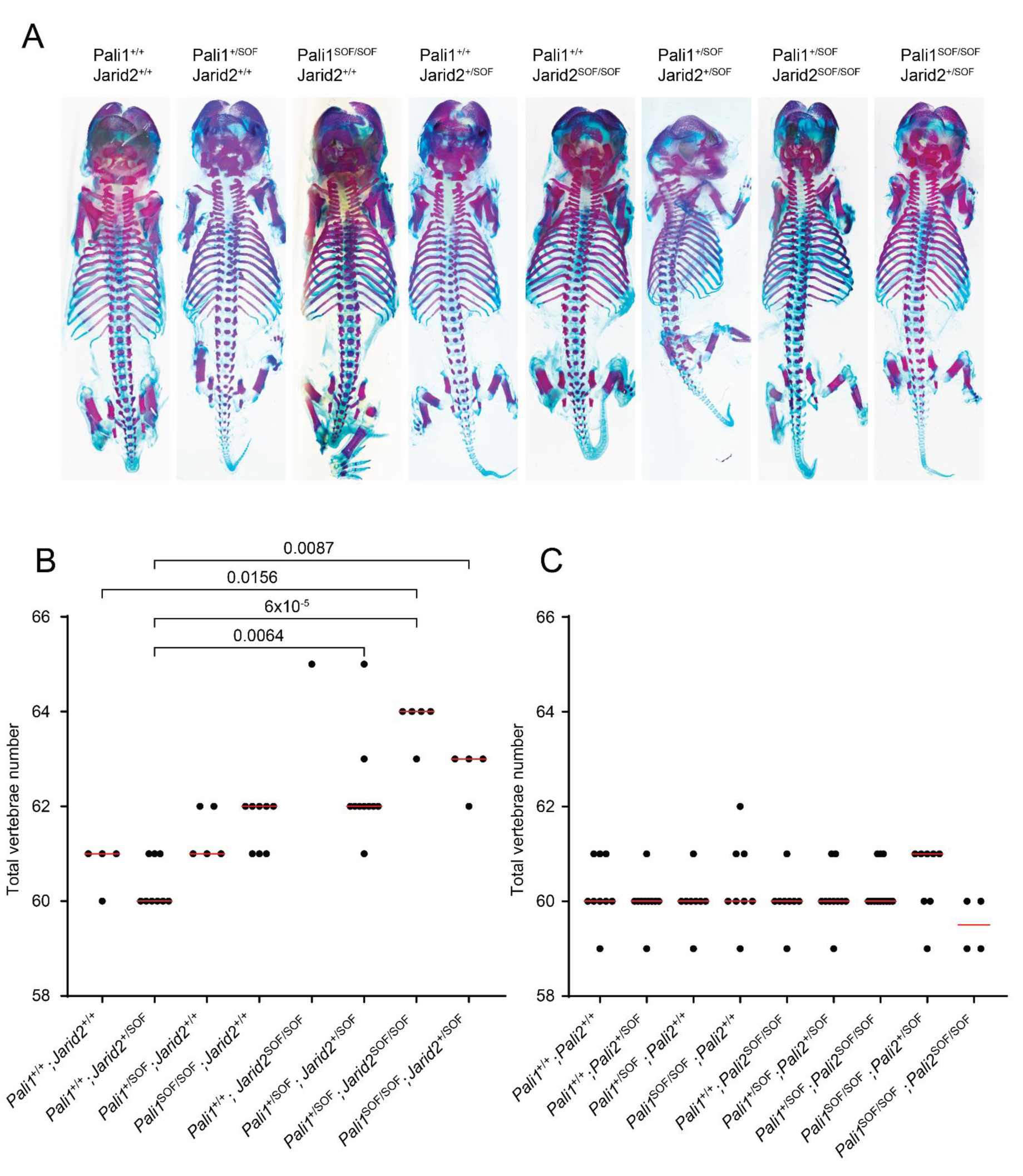
The synergy between allosteric defective JARID2 and PALI1 during mouse embryogenesis. (A) The photos shown in this figure correspond to the cropped photos shown in Figure 3C. An exception is the *Pali1^+/SOF^;Jarid2^+/SOF^* genotype, which was photographed using a microscopic lens limited only to the area of interest in Figure 3C, whereas a photograph of the complete embryo that was recorded separately is shown here, positioned similarly. Genotypes are indicated. B. Total vertebral number for the embryos that were part of the experiment and are shown in Figure 3C. Each dot on the graph represents the total vertebral number for one embryo. Only embryos for which all vertebrae could be confidently counted were included on this plot. The median for each genotype is shown by a red line. Genotypes are indicated. A Kruskal-Wallis test with Dunn’s test for multiple comparisons was used to test for significance between all combinations of genotypes and p values are presented only where differences are statistically significant. C. Same as (B), except that the embryos were part of the experiment shown in Supplementary Figure 2C.

**Supplementary Figure 4:**
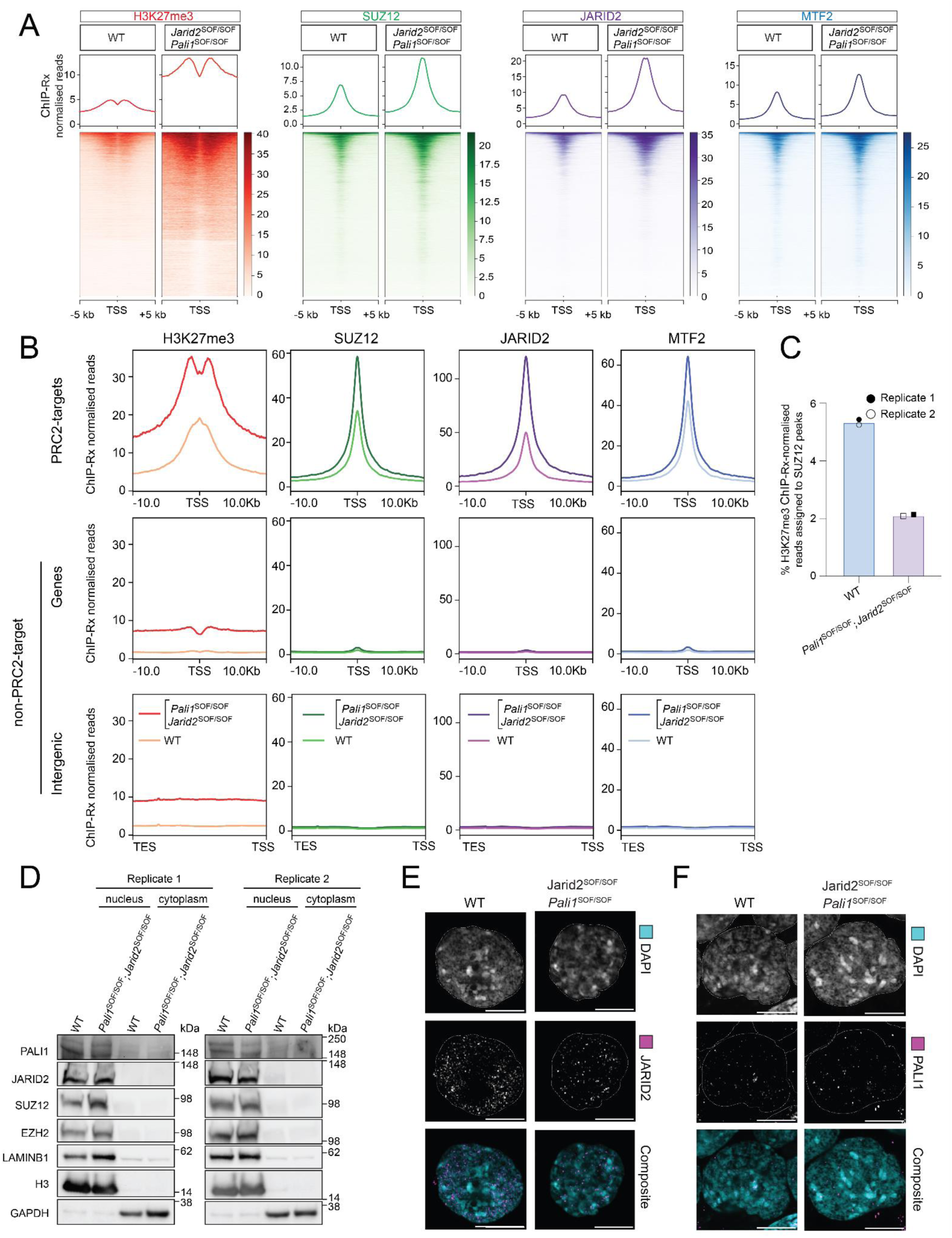
Allosteric defective JARID2 and PALI1 lead to a genome wide increase of H3K27me3 in mESC. A. Second replicate of the ChIP-Rx shown in Figure 4D, where heatmaps are centred on all transcription start sites. B. Same data set as in (A), where average ChIP-Rx signal profiles are shown over all transcription start sites at PRC2-target genes, non-PRC2-target genes, or intergenic regions as indicated. C. Bar chart representing the percentage of ChIP-Rx-normalised H3K27me3 reads that aligned to genomic regions within SUZ12 peaks in the wild-type and mutant cell lines within two independent ChIP-Rx replicates. D. Western blots on the nuclear or cytoplasmic fractions of the wild-type or double mutant *Pali1^SOF/SOF^;Jarid2^/SOF/SOF^* cells, using antibodies as specified. E. Representative fluorescence confocal microscopy images of wild-type or double mutant *Pali1^SOF/SOF^;Jarid2^SOF/SOF^* cells as indicated. Channels for each stain are shown from top to bottom: DAPI (DNA staining in cyan), anti-JARID2 antibody (magenta) and merged composite image. All scale bars represent 10 µm, and outlines delineate the extent of the nuclei. F. Representative fluorescence confocal microscopy images as in (E), except that anti-PALI1 antibodies were used. Channels for each stain are shown from top to bottom: DAPI (DNA staining in cyan), anti-PALI1 antibody (magenta) and merged composite image.

**Supplementary Figure 5:**
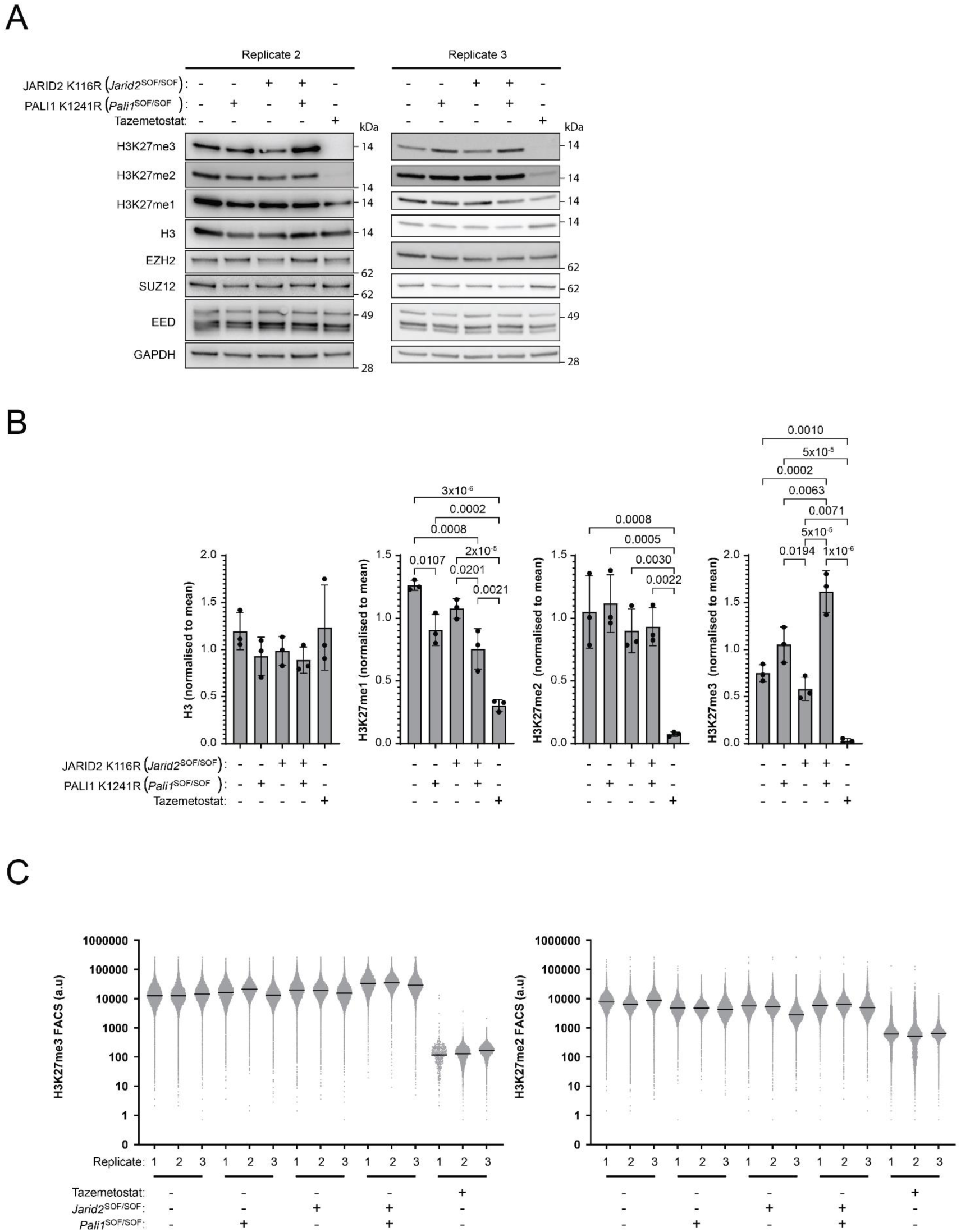
Allosteric defective JARID2 and PALI1 lead to increased global H3K27me3 in mESC. A. Two more western blot replicates corresponding to Figure 4C. B. Bar charts representing the mean densitometry values of all three western blot replicates shown in (A) and in Figure 4C. Shown are densitometry values that were normalised to the average densitometry values of all cell lines in the experiment, excluding the Tazemetostat treated sample. C. Flow cytometry data corresponding to Figure 4B. Indicated are individual fluorescence values for each cell that participated in the experiments. Cell lines and Tazemetostat treatments are indicated. The medians that are presented in Figure 4B were derived from this analysis and are presented here with horizontal black lines. Each replicate is plotted separately, and three independent replicates of each sample are grouped together.

**Supplementary Figure 6:**
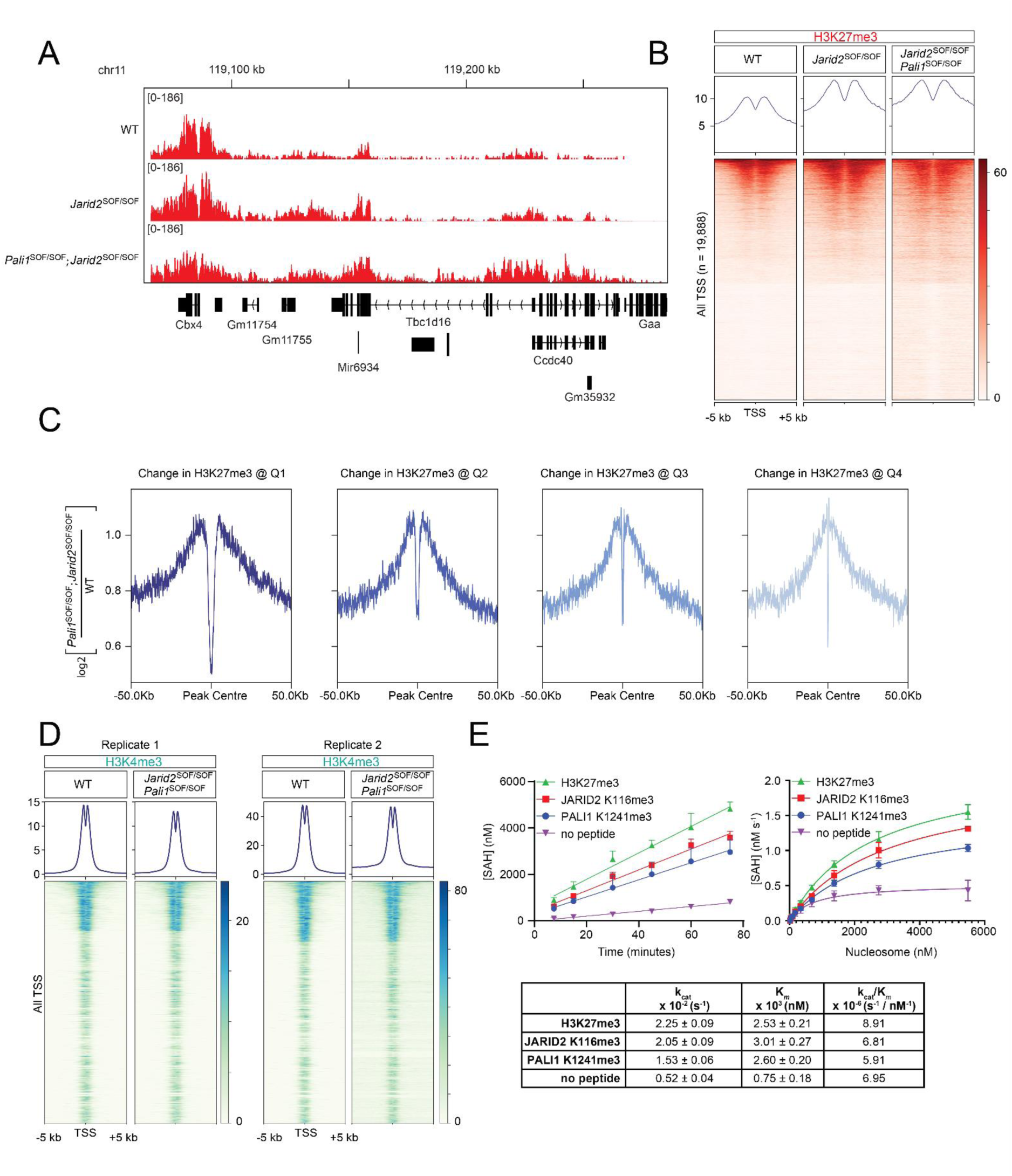
The JARID2 K116R mutant exhibits increased H3K27me3 in cells. A. Gene track over a representative locus showing H3K27me3 ChIP-Rx signals from cell lines as indicated. B. Heatmaps corresponding to the ChIP-Rx shown in (A), centred on all transcription start sites. C. Relative enrichment profiles of the ChIP-Rx data from double mutant *Pali1^SOF/SOF^;Jarid2^/SOF/SOF^* cells divided by the ChIP-Rx data from the wild-type cells, and centered over H3K27me3 peaks that were called based on the WT cells and were splitted into four quartiles, Q1 to Q4, as indicated. In this analysis, Q1 and Q4 represent the top and bottom 25% H3K27me3 peaks in the wild-type cells, respectively, based on peaks width. The values represent the ratio between H3K27me3 in the mutant normalised to the wild type. D. Quantitative ChIP-seq (ChIP-Rx) of H3K4me3 in cell lines as indicated. E. Histone-methyltransferase assays were carried out for PRC2 in the presence of H3K27me3, JARID2 K116me3, or PALI1 K1241me3 allosteric effector peptides, as indicated. Left: a progress curve was carried out using 100 nM PRC2 in the presence or absence of 100 µM peptides, as indicated, and 5.55 µM mononucleosomes in reaction buffer of 50 mM Tris pH 8.0 at 30C, 100 mM KCl, 0.5 mM MgCl2, 0.1% Tween 20, 5 mM DTT and 25 µM SAM. The SAH concentration was measured at 7.5, 15, 30, 45, 60, and 75 minutes. Right: a substrate titration assay in the presence of 100 nM PRC2. Each point represents the mean rate of SAH production at a given mononucleosome concentration, titrated from 0 to 5.55 µM, and incubated for 120 minutes in reaction buffer as used for the progress curve (left), with 100 µM peptides as indicated. The trendlines represent a non-linear regression. Each data point represents the mean SAH concentration (left) or the rate of SAH production (right), as determined by luminescence against a standard curve. Error bars represent the standard deviation over three independent replicates carried out on different days. *k*_cat_, *K*_M_, and the catalytic efficiency (*k*_cat_/*K*_M_) values are indicated with standard errors in the table underneath the plots.

**Supplementary Figure 7:**
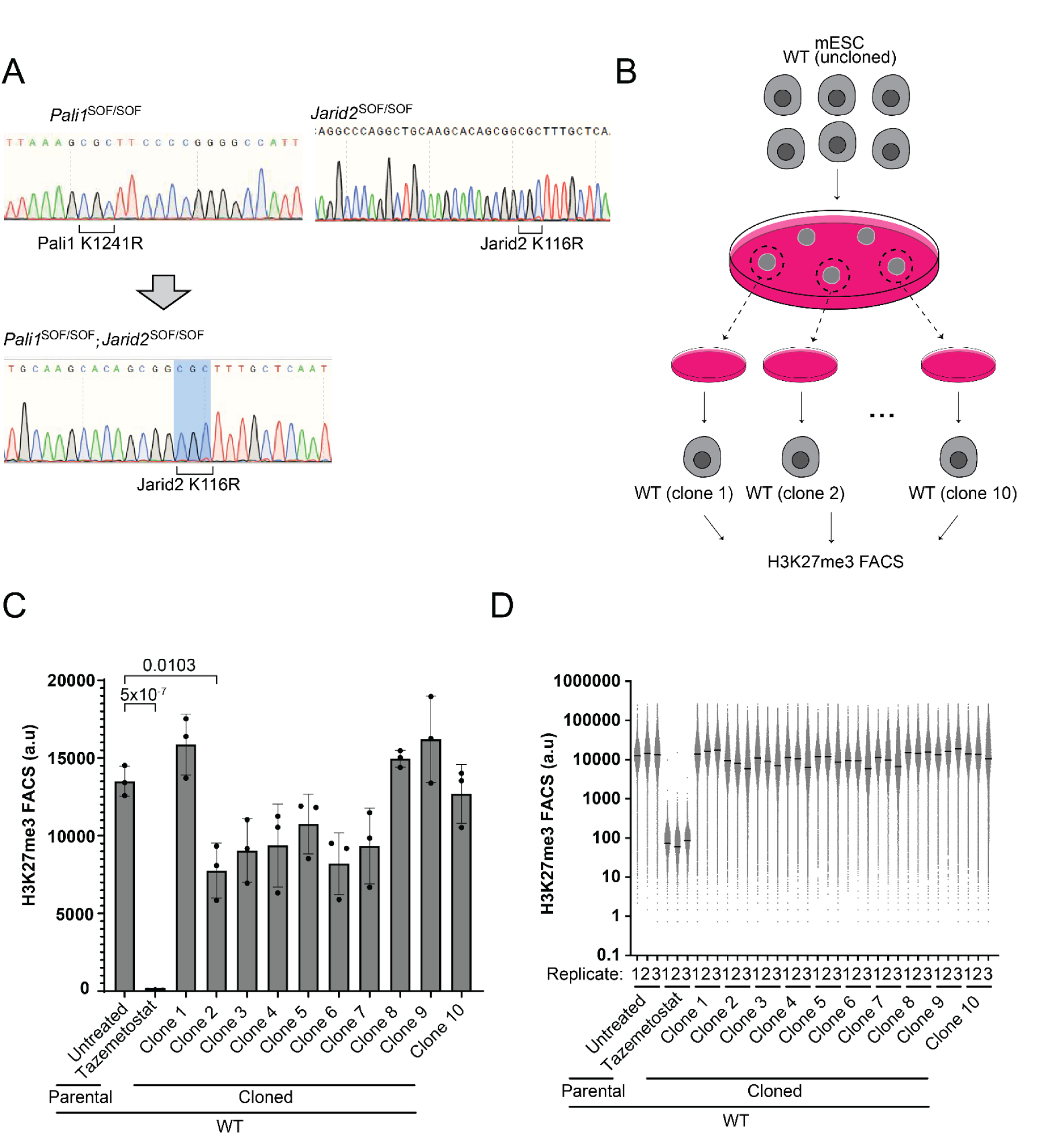
Clonal selection of wild-type mESCs is insufficient to cause a global gain of H3K27me3. A. Sanger sequencing traces confirming the correct knock-in homozygous mutant mESC lines generated in this study. The codon corresponding to the separation of function mutation is annotated below. B. A schematic diagram showing how the cells were treated to assess for a clonal effect: The parental wild-type mESCs were seeded at a low density on a culture dish, and incubated until colonies grew from the individual cells. Ten colonies were picked and transferred into separate dishes. The resulting ten clonal wild-type cell lines were all expanded and then assayed for global H3K27me3 by flow cytometry. C. Flow cytometry of the ten clonal wild-type cells (see scheme in (B)) stained with anti-H3K27me3 antibodies. The “parental” sample represents the same parental wild-type cell line used in other experiments and also for clonal selection here. Tazemetostat was used as a negative control. The bar charts show the mean of the median fluorescence signal and error bars represent standard error across three independent biological replicates that were harvested on different days. Statistical significance was calculated using a one-way ANOVA with a post-hoc Tukey’s test to correct for multiple comparisons. Only comparisons involving the uncloned cells are shown. The p-values of non-significant comparisons are not shown. D. Scatter plots of the H3K27me3 flow cytometry values measured from each cell within the cell lines shown in (B). This data was used to calculate the median values (marked by a line) that are presented in (C). X-axis labels are as in (C), except that three independent replicates are presented side-by-side for each of the wild-type clonal cell lines.

**Supplementary Figure 8:**
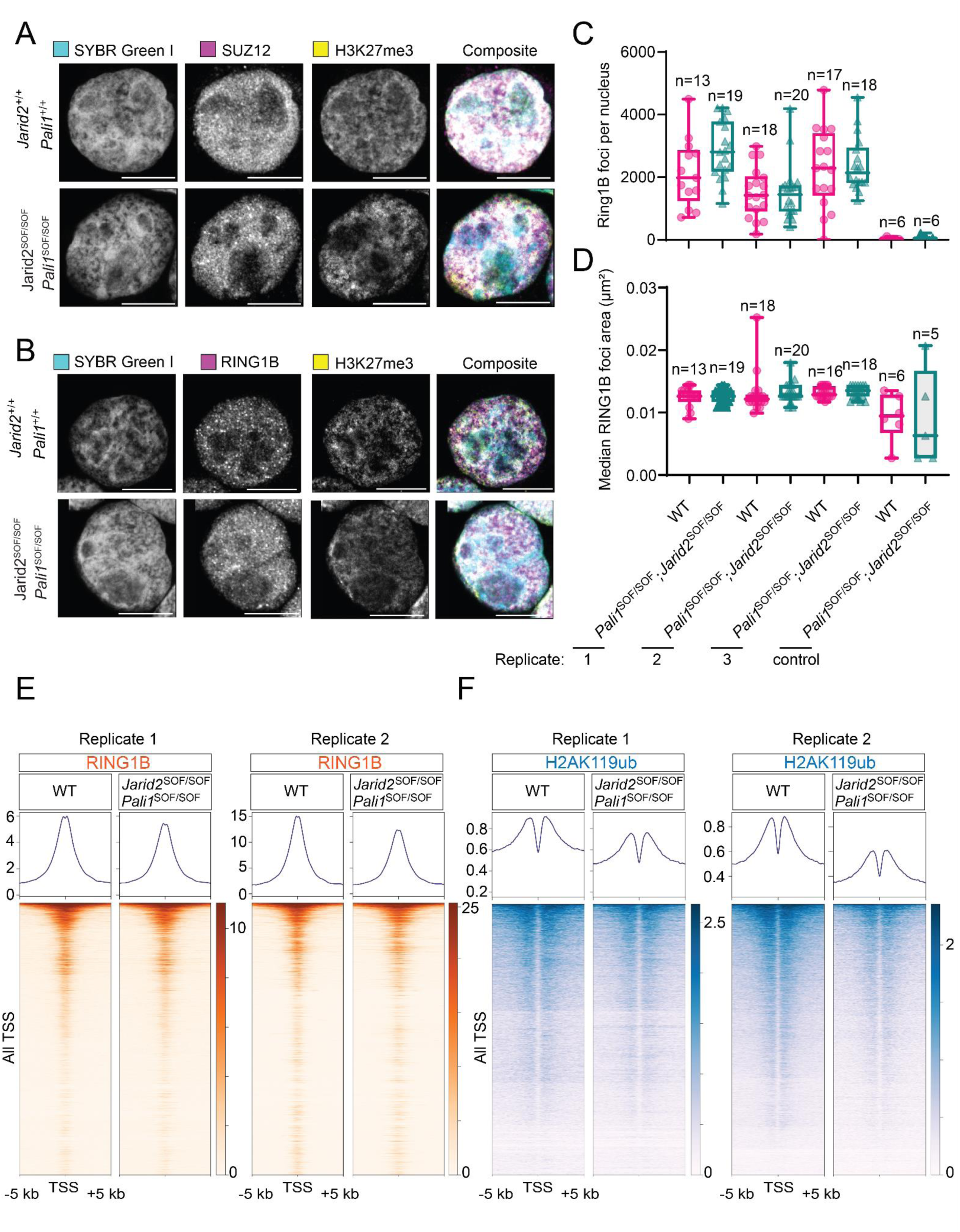
Defective H3K27me3 mimicry does not substantially change the chromatin occupancy of RING1B. A. Representative stimulated emission depletion (STED) super resolution immunofluorescence microscopy images of wild-type or double mutant *Pali1^SOF/SOF^;Jarid2^/SOF/SOF^* cells, as indicated. Channels for each stain are shown from left to right: SYBR Green (DNA staining, in cyan), anti-SUZ12 antibody (magenta), anti-H3K27me3 antibody (yellow), and merged composite image. Scale bars represent 10 µm. B. Representative fluorescence microscopy images as in (A), except that an anti-RING1B antibody (magenta) was used instead of anti-SUZ12 antibody. C. Box plots showing the number of RING1B foci in the nuclei of the wild-type (pink) or double mutant (green) cells, based on the imaging in (B). The number of cells assayed is given as “n”, and data from three independent replicates that were carried out in three different days is shown. The boxes extend from the 25^th^ to 75^th^ percentiles, the middle line represents the median, and the whiskers represent the minimum and maximum values. The condition labelled “control” refers to cells that were treated as the cells above, except without primary antibody. D. As in (C), except that the plots represent the median area of the foci instead of the number of foci. E. ChIP-Rx using anti-RING1B antibodies in the wild-type or the double mutant cells, as indicated. Heatmaps are centred on all transcription start sites. F. As in (E), except that anti-H2AK119ub antibodies were used.

**Supplementary Figure 9:**
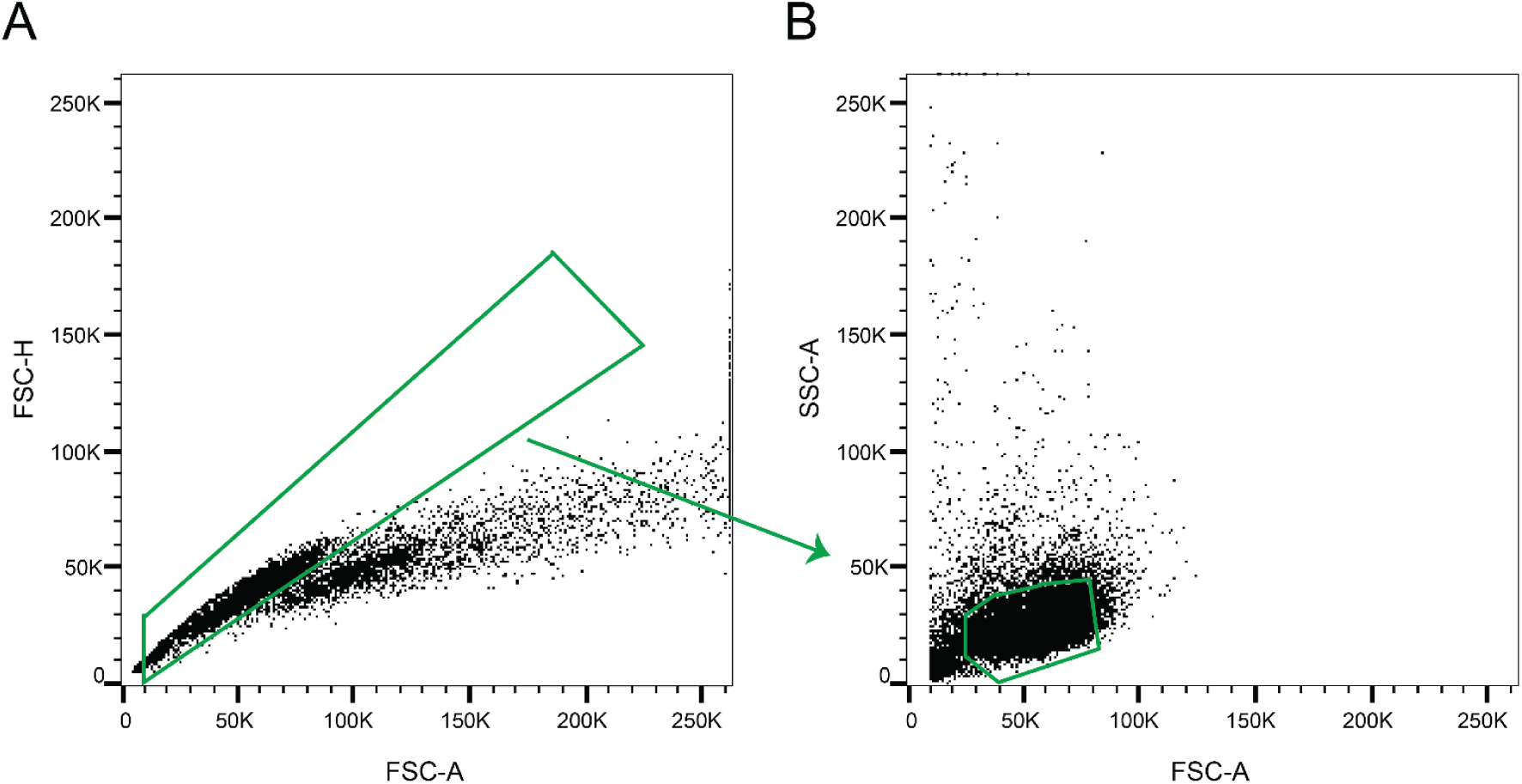
Gating strategy used for flow cytometry analysis. Representative scatter plots showing the gating strategy used for measuring global H3K27me2/3 by flow cytometry. A gate was used to include only the single cells (A), and another gate to include the intact mESCs (B). Only the events that fell within both gates were considered when measuring the median H3K27me2 or H3K27me3 for that sample.

**Supplementary Data Table 1: Embryonic skeleton phenotypes.** A spreadsheet showing the phenotype along the axial skeleton of each embryo included in the experiments shown in Figures 1G and 3C and Figures S1B, S2B,C and S3. Scoring of the skeletons resulting from unique crossings of genotypes are provided in separate tabs. A key is shown below the table in each tab.

**Supplementary Data Table 2: Differential expression analysis in mouse embryonic tailbuds using RNA-seq.** A spreadsheet containing DESeq2 differential expression analyses of RNA-seq data generated using RNA that was extracted from mouse embryonic tailbud tissue samples. Provided in separate tabs are separate differential expression analyses corresponding to each of the comparisons shown in the heatmap in Figure 2C. The data in each tab is the direct output of the DESeq2 results function, with no further processing or sorting.

**Supplementary Movie 1: Wild-type E12.5 embryo immunostained for SOX9.** A movie showing the complete light sheet fluorescence microscopy 3D reconstruction in Figure 1F. First, the complete signal is shown without removing any non-relevant regions. Next, the non-relevant signal is removed, as in the second row of Figure 1F. Next, a rendering of both of the pairs of ribs are imposed over the previous model, and finally only the side used to generate the bottom row in Figure 1F is shown.

**Supplementary Movie 2: *Jarid2^+/SOF^* E12.5 embryo immunostained for SOX9.** As Supplementary Movie 1, except that a *Jarid2^+/SOF^* E12.5 embryo was imaged.

Supplementary Movie 3: *Jarid2^SOF/SOF^* E12.5 embryo immunostained for SOX9. As Supplementary Movie 1, except that a *Jarid2^SOF/SOF^* E12.5 embryo was imaged.

## Methods

### Amino acid numbering system

For compatibility with previous works, the allosteric effector lysines in mouse cells and animals are named based on the human numbering system. Accordingly, references to PALI1 K1241, PALI2 K1558 or JARID2 K116 (PALI1 K1393, PALI2 K1543 or JARID2 K116, respectively, in mouse) are made in this way, using the human numbering system, regardless of if these amino acids are in the human or mouse proteins. The corresponding sequences of the mouse and human proteins are shown in Figure 1B, where the human and mouse sequences were aligned using EMBOSS needle ^71,72^.

### Mice genotyping and ethical approval

All mice used in this study were maintained on a C57BL/6J background. The genetic background was confirmed by Transnetyx using miniMUGA. Mouse lines possessing mutations in multiple genes were generated by crossing the relevant single mutant lines. Genotyping was performed by Transnetyx using real-time PCR. All animal breeding and experimentation was conducted following the Australian National Health and Medical Research Council Code of Practice for the Care and Use of Animals for Scientific Purposes guidelines for housing and care of laboratory animals and performed in accordance with Institutional regulations after pertinent review and approval by Monash University Animal Ethics Committee under project numbers 36532 and 26125. Mice were bred and housed in specific-pathogen-free (SPF) conditions at the Monash Animal Research Platform (MARP) and Animal Research Facility (ARL) at Monash University.

### Knock-in of point mutations to mice

PALI1 K1241R, PALI2 K1558R and JARID2 K116R were knocked-in using CRISPR/Cas with homology-directed repair, using sgRNAs that were designed to target the following sequences: TCCTAGCAGGAATGGCTCCA, AAGGAGAAGAGACTGCAAATT and GGCCCAGGCTGCAAGCACAA, respectively. To avoid embryonic lethality resulting from homozygous knock-in of the mutation, two repair templates were used for each genome modification process: one that contains an MspA1I restriction site but without amino acid change (Repair Template 1), and one that contains MspA1I and HaeII restriction sites, and also the amino acid change (Repair Template 2).

The appropriate sgRNA and Cas protein were incubated with the guide RNA to form a ribonucleoprotein (RNP) complex. Cas9 used the PALI1 K1241R and JARID2 K116R knock-in and Cas12a used for the PALI2 K1558R knock-in. The RNP complex and repair template were then electroporated or microinjected into C57BL/6J zygotes at the pronuclei stage and transferred into the uterus of pseudo pregnant F1 females. The following conditions were used for each gene:

PALI1 K1241 knock-in: Repair Template 1 sequence was CCTTCCACCATGTACCCTAGTTCACTACAGGCAGAACGCTTGAAAAAACATTTAAAGAAATTCCCCGGGGCCATTCCTGCTAGGAATAATTGGAAGACACAGAAGCTATGGGCTAAACTACGAGAGAATCCTGA. Repair Template 2 was AGTTTCCTCCTTCCACCATGTACCCTAGTTCACTACAGGCAGAACGCTTGAAAAAACATTTAAAGCGCTTCCCCGGGGCCATTCCTGCTAGGAATAATTGGAAGACACAGAAGCTATGGGCTAAACTACGAGAGAATCCTGA. Delivery was done using the microinjection of Cas9 nuclease (IDT Alt-R S.p. Cas9 Nuclease V3; 10 ng/mL), sgRNA (10 ng/mL) and repair templates (30 ng/mL).

PALI2 K1558R knock-in: Repair Template 1 was TTTAAGAGTGGTGAGCCAACCATTTTATGTAGTATATGAAATTTCCCAGCTGGTTTTGAAGGAGAGCTCACTGCAAATTTTTTAAAGTGCTTCCTGAAGGGGCTGGCATTAAATCCACAGCATTTCACCCCTAACTTAATGCCCT. Repair Template 2 was TTTAAGAGTGGTGAGCCAACCATTTTATGTAGTATATGAAATTTCCCAGCTGGTTTTGAAGGAGAGCTCACTGCAAAACGTTTAAAGTGCTTCCTGAAGGGGCTGGCATTAAATCCACAGCATTTCACCCCTAACTTAATGCCCT. Delivery was done using the microinjection of Cas12a nuclease (Alt-R A.s. Cas12a (Cpf1) Ultra; 100 ng/mL), crRNA (100 ng/mL) and repair templates (50 ng/mL).

JARID2 K116R knock-in: Repair template 1 was CTCCAGTCTCTGCTTAATTTAGTTTTACATTTCTTTTGTTGTTGCAGGCCCAGGCTGCAAGCACAGCGGAAGTTTGCTCAATCTCAGCCGAATAGTCCCAGCACAACTCCAGTGAAGATAGTGGAGCCTTTG. Repair template 2 was CTCCAGTCTCTGCTTAATTTAGTTTTACATTTCTTTTGTTGTTGCAGGCCCAGGCTGCAAGCACAGCGGCGCTTTGCTCAATCTCAGCCGAATAGTCCCAGCACAACTCCAGTGAAGATAGTGGAGCCTTTGCTACC. Delivery was done using the microinjection of Cas9 nuclease (IDT Alt-R S.p. Cas9 Nuclease V3; 10 ng/mL), sgRNA (10 ng/mL) and repair templates (30 ng/mL) or electroporation of Cas9 nuclease (3.8 mM), sgRNA (4.2 mM) and repair templates (10 mM). For all knock-ins, the genotypes of the founder and F1 mice were confirmed by sanger sequencing (Figure S2A).

### Analysis of mendelian inheritance

Mice possessing the desired genotypes were crossed. The resulting pups that survived to 21 days postnatal were genotyped, and the frequency of each genotype among them was tallied. The ‘expected’ number of mice represents the number of pups of a given genotype expected to survive if all the possible genotypes resulting from the given crossing are viable. This was calculated using Punnett squares of monohybrid and dihybrid crosses in cases of one or two mutant genes, respectively. Chi-square tests were conducted using Excel for the pups resulting from the crossings to test for statistical significance between the observed and expected ratios of genotypes.

### Skeleton preparation and imaging

Mice heterozygous for each of the mutations of interest were time-mated, and embryos were harvested at embryonic day 14.5 or 18.5 (E14.5 or E18.5, respectively).

E18.5 embryos were prepared as described previously ^73^. Specifically, the E18.5 embryos were skinned, then incubated in 95% (v/v) ethanol for 2 days, followed by 2 days in acetone. The embryos were then stained for 1-2 days in a solution containing 150 µg/mL alcian blue (Sigma 05500), 50 µg/mL alizarin red (Sigma A5533), 5% (v/v) glacial acetic acid and 60% (v/v) ethanol. The embryos were then cleared in 1% (w/v) KOH for 3-5 days, then transferred through a graded series of solutions in which a gradually increasing proportion of the KOH was replaced with glycerol (25%, 50%, 75% then 100% glycerol), with one day between each solution change. All incubations took place at room temperature with rocking.

E14.5 embryos were stained as previously described ^74^ with some minor changes. Specifically, the E14.5 embryos were incubated while rocking in 70% (v/v) ethanol at 4°C overnight, then at room temperature in 95% (v/v) ethanol for 1 hour, followed by acetone overnight. The embryos were stained in a solution containing 0.03% (w/v) alcian blue, 80% (v/v) ethanol and 20% (v/v) acetic acid, for 4 hours. The embryos were cleared over 2-3 days in 0.5% (w/v) KOH, then transferred through a graded series of solutions in which a gradually increasing proportion of the KOH was replaced with glycerol (25%, 50%, 75% then 100% glycerol), with one day between each solution change. All incubations took place at room temperature with rocking.

Images of E18.5 embryos were acquired using a Vision Dynamic BK Lab Imaging System equipped with a Canon 5D Mark II with a 100 mm macro lens, or 7D Mark II with Mitutoyo 5x microscope lens. Z-stacked images were captured at multiple focal planes through the embryo, then combined to create a composite photograph using Zerene Stacker. Images of E14.5 embryos were taken with a ZEISS Stemi 508 microscope equipped with a ZEISS Axiocam 208 color camera. The images were set to black and white, and brightness and colour levels for each image were adjusted in Photoshop separately.

### RNA sequencing of mouse embryonic tailbuds

Mice heterozygous for the JARID2 K116R mutation (*Jarid2^SOF/+^)* were time-mated, embryos harvested at approximately embryonic day 9.5 (E9.5), and the yolk sac was genotyped by Transnetyx using real-time PCR. All embryos resulted from crossing *Jarid2^SOF/+^* animals with the exception of wild-type embryos of 12 to 14 somites that were generated from a wild type intercross that were collected approximately 0.5 days earlier (E9.0, i.e. 12 to 14 somites), since the JARID2 K116R mutant embryos exhibited a small developmental delay compared to their littermate wild-type embryos. This was done to ensure that mutant and wild-type embryos were assayed at the same somite stage of development, allowing a direct comparison of *Hox* gene expression levels which are known to be highly dynamic in the tailbud at these stages of development ^75^.

Embryos were dissected out of uterine horns in cold PBS. Immediately after harvesting each embryo, the tailbud tissue was removed, placed in 350μL lysis buffer (buffer RLT) from an RNeasy mini kit (Qiagen 74106), and was then snap frozen in dry ice and stored at -80°C. The remainder of the embryo was fixed overnight in 4% paraformaldehyde (PFA) solution in PBS (Santa Cruz sc-281692) at 4°C with rocking. After fixing, the embryos were washed in PBT (PBS with 1% Tween-20) then dehydrated by transferring through a series of PBT-methanol solutions, each containing a higher concentration of methanol (25%, 50%, 75%, 100% methanol) for 5 minutes each, then stored at -20°C in 100% methanol until further processing.

In-situ hybridisation was used to stain the somites in the fixed embryos, using an RNA probe against *Uncx4.1*. After staining, the number of somites were counted and recorded for each embryo.

The solution containing the tailbud tissue was passed through a Qiashredder column (Qiagen 79654), and RNA then was extracted and purified using the RNeasy mini kit according to the manufacturers instructions, including DNase digestion. The purified RNA was dried overnight at room temperature in a biosafety hood in RNA stabilisation tubes (GenTegra GTR5025-S) before shipment at room temperature to the sequencing service provider (Azenta). RNA was subjected to SMRT-based library preparation and Illumina sequencing, aiming for 30 million paired-end reads per library at a 2×150 bp format (Azenta).

### Bioinformatic analysis of RNA sequencing data

All fastq files were trimmed using Trim Galore (version 0.5.0), then transcripts were quantified using Salmon (version 1.5.2) ^76^ against cDNA sequences from the mm10 build indexed with Salmon. Gene-level count matrices for differential expression analysis were generated from the Salmon transcript quantification output files (quant.sf files), with experimental groups (i.e. genotype and somite number) annotated using a CSV annotation table (named sample_sheet.csv) that contained a “sample” column containing the names of the quant files for each embryo, and a “condition” column containing the corresponding experimental group that each embryo belongs to. Transcripts were mapped to genes using the TxDb.Mmusculus.UCSC.mm10.knownGene annotation package. Transcript-level abundance estimates were then summarised to the gene level with tximport, and passed to the DESeq function from DESeq2 (version1.46.0) ^77^. For differential expression analyses, the resulting DESeq2 function was passed to the results function with default settings. Genes were labelled by mapping their Entrez ID to the corresponding mouse gene symbol, using the mapIds function in the org.Mm.eg.db package (version 3.20.0). The object containing differential expression analysis with gene names was then written to a CSV file.

To create heatmaps of Hox genes expression, a regularised log transformation was applied to the DESeq2 object using the rlog function from DESeq2. Gene symbols were mapped to Entrez IDs through the same process as in the differential expression analysis above. The rlog transformed dataset was then filtered to only include the gene symbols corresponding to the genes to be plotted. To calculate changes in gene expression between multiple groups of embryos in the same plot, embryos were partitioned into pairs of groups (i.e. group A and group B) based on their number of somites and genotype (e.g. wild type with 12 somites, wild type with 13 somites, etc.), as defined within the “sample_sheet.csv” file that is described above, in the “condition” column. For each comparison between two groups of embryos, the change in expression for a given gene was calculated as the difference between the mean rlog expression in group B and the mean rlog expression in group A (i.e. mean(rlog(group B)) - mean(rlog(group A)). The resulting matrix was visualized using the pheatmap package (version 1.0.12).

### 3D light sheet immunofluorescence microscopy of whole embryos

To visualise the developing skeleton, *in toto* immunofluorescence was performed as described previously ^78^. E12.5 embryos were dissected out of uterine horns in cold PBS and fixed overnight in PBS with 4% PFA at 4°C with rocking. The following day, embryos were washed in PBS then dehydrated by transferring through a series of PBS-methanol solutions, each containing a higher concentration of methanol (20%, 40%, 60%, 80%, 100% methanol) for one hour each, then frozen in 100% methanol until further processing. Prior to staining, embryos were bleached in methanol with 7.5% H2O2 under white light at 4°C with rocking, and rehydrated by transferring through a series of PBS-methanol solutions, each containing a lower concentration of methanol (80%, 60%, 40%, 20%, 0% methanol) for one hour each.

The rehydrated embryos were permeabilised in PBS with 0.2% Triton X-100, 20% DMSO and 2.3% glycine, then blocked in PBSGT (PBS with 0.2% gelatin and 0.5% Triton X-100). Permeabilisation and blocking were performed at 37°C with rocking for 48 hours.

The embryos were incubated with anti-SOX9 (Merck AB5535, 1:100) and anti-TUBB3 (BioLegend 801201, 1:500) primary antibodies diluted in PBSGT at 37°C in the dark with rocking for at least 5 days. After the primary antibodies were removed, the embryos were washed at least four times in PBSGT for one hour per wash. The embryos were then incubated with anti-rabbit (Thermo Scientific A-31572, 1:500) and anti-mouse (Thermo Scientific A-21241, 1:500) secondary antibodies for at least four days at 37°C in the dark with rocking. After the secondary antibodies were removed, the embryos were washed at least four times in PBSGT for one hour per wash.

Next, the embryos were dehydrated again by transferring through a series of PBS-methanol solutions, each containing a higher concentration of methanol (20%, 40%, 60%, 80%, 100% methanol) for one hour each at room temperature in the dark, rotating. This was followed by another wash in methanol for one hour, then again overnight. Clearing was performed with a 2:1 ratio of dichloromethane (DCM) to methanol for 3 hours in the dark, rotating, followed by 100% DCM for one hour, and finally submerged into dibenzyl ether (DBE) without further agitation to avoid excessive oxidation. Embryos were imaged within one week of clearing.

Images were acquired on an Ultramicroscope 2 (La Vision, Miltenyi Biotec) equipped with a 2X objective and a 1.6X magnification in DBE. Single side illumination, with 3 convergent light sheets set at a width of 80% and a numerical aperture of 0.134 Nyquist was used to scan the sample. Dynamic focus was used across the X axis, with the proprietary blend algorithm in Imspector Pro 5.0 (La Vision, Miltenyi Biotec).

Image processing was performed in Imaris 10.2 (Bitplane, Oxford Instruments). The SOX9 signal for the rib cage and spine was isolated by manually subtracting signal from other non-relevant SOX9-expressing tissues (limbs, heart, skin, gonadal ridges) using the surface function in Imaris. 3D rendering of the ribs was generated using the surface function in Imaris. Images were acquired using the Snapshot function of Imaris, and the brightness was adjusted in ImageJ. Movies were generated using the Animation menu of Imaris.

### Cell culture

Bruce-4 mouse embryonic stem cells were cultured at 37 °C with 5% CO2 in DMEM growth medium, supplemented with 15% FBS (Scientifix FBSAU-2007A), 1% penicillin-streptomycin (Thermo Scientific 15140122), MEM Non-Essential Amino Acids Solution (Thermo Scientific 11140050), Glutamax™ Supplement (Thermo Scientific 35050061), 0.1 mM 2-Mercaptoethanol (Merck M3148) and 1:1000 homemade LIF. The cells were grown on gelatinised plates, passed every 2-3 days, and the media was replaced daily. Bruce-4 cells (derived from C57BL/6J-Thy1.1 strain, kindly gifted to Monash Genome Modification Platform by Colin Stewart) were authenticated by ATCC using short tandem repeat (STR) analysis. Cells were tested periodically for mycoplasma contamination using PCR ^79^. In experiments where H3K27me3 level was assessed using flow cytometry and immunoblotting, a Tazemetostat-containing control was used to identify the background level detected by the H3K27me3 antibodies in the absence of H3K27me3 in the cells.

### Knock-in of point mutations to mESCs

PALI1 K1241R, PALI2 K1558R and JARID2 K116R were knocked-in using CRISPR/Cas with homology-directed repair, using the sgRNAs that were designed for the following targets: TCCTAGCAGGAATGGCTCCA, AAGGAGAAGAGACTGCAAATT and GGCCCAGGCTGCAAGCACAA, respectively. Lipofectamine CRISPRMAX was used to transfect mESCs with the appropriate sgRNA, Cas9 protein fused with GFP and repair template with the same sequence that was indicated above, for mouse knock-in. Clonal cells were selected using flow cytometry to sort individual GFP-positive cells. In all cases, clonal cells were expanded and screened for the desired genotype using PCR and restriction enzyme digestion. Positive clones were validated by sanger sequencing (Figure S7A). Cell lines were occasionally genotyped by Transnetyx using real-time PCR.

### Detection of H3K27me2/3 using flow cytometry

For a negative control, wild-type cells were grown in 10 µM Tazemetostat (Selleckchem S7128) for 7 days. Other cell lines were cultured without Tazemetostat. To collect cells, the culture media was removed, replaced with trypsin (Thermo Scientific 25200-056) solution and cells were incubated in trypsin at room temperature for <5 minutes. The detached cells were resuspended in growth media and collected by centrifugation for 5 minutes at 300 g. The cell pellet was then resuspended in 4% paraformaldehyde solution in PBS (Santa Cruz sc-281692) and incubated for 15 minutes at room temperature to fix the cells. The cells were then washed twice in PBSTX (PBS with 0.1% (v/v) Triton X-100 (Sigma X100)), and resuspended in PBSTX to be stored for up to a month at 4 °C before further processing.

For the antibody staining, 2×10^5^ cells per sample were resuspended in blocking buffer (PBS with 0.1% (v/v) Triton X-100, 1% (w/v) BSA (Sigma 10735078001) and 5% (v/v) FBS) and blocked for 15 minutes.

The cells were centrifuged and resuspended in blocking buffer with H3K27me2 (Abcam ab24684, 1:5000) or H3K27me3 antibody (Cell Signalling #9733S, 1:500). After 1 hour of incubation, the cells were then washed with PBSTX, centrifuged at 1000 g for 2-5 minutes and the pellet resuspended in blocking buffer with secondary antibody (Thermo Scientific A32795, 1:15000) and 1 µg/mL DAPI (Sigma, D9542-1MG) for 45 minutes in the dark. The cells were then washed with PBSTX, centrifuged at 1000 g for 2-5 minutes, and resuspended in PBS with 10% FBS and 615 µM EDTA, and ran through a cell strainer (Falcon #352235). The cells were then analysed on a BD LSRFortessa™ X-20 Cell Analyzer, using the R670-A detector.

Three independent biological replicates were initiated on three different days and were carried out as described above. The data was gated for intact single cells (Figure S9) using FlowJo and plotted using GraphPad Prism.

### Immunoblotting

To collect cells, the culture media was removed, replaced with trypsin and incubated in the trypsin solution for <5 minutes at room temperature. The detached cells were resuspended in growth media and collected by centrifugation for 5 minutes at 300 g. The cells were then washed 3 times in PBS and lysed in Laemmli buffer (1% (v/v) SDS, 12.5% (v/v) glycerol, 35 mM Tris pH 7.5 at 25°C, 0.01% (w/v) bromophenol blue, 5 mM MgCl2, 1% (v/v) 2-mercaptoethanol) and 25 U/mL Benzonase (Merck #70746) to extract whole cell lysate. The samples were then heated to 95 °C for 5-10 minutes before they were loaded on a gel. Where nuclear fractionation was used, after washing with PBS cells were instead resuspended in cytoplasmic extraction buffer (20 mM Tris pH 7.5 at 25 °C, 0.1 mM EDTA, 2 mM MgCl2, 20 mM BME and protease inhibitor cocktail (Sigma #4693132001)). The cells were incubated for 2 minutes at room temperature, then 10 minutes on ice before adding NP-40 to a concentration of 1% (v/v) and mixing. Samples were centrifuged at 4 °C and 500 g for 3 min and the supernatant was kept as the cytoplasmic fraction and 25 U/mL Benzonase was added. The nuclear fraction was washed in cytoplasmic extraction buffer with 1% (v/v) NP-40 twice, by centrifugation at 4 °C at 500 g for 3 min. To lyse nuclei, the pellet was resuspended in 500µL of high salt lysis buffer (50 mM Tris pH 8 at 25 °C, 1 mM EDTA, 0.5% (v/v) NP-40, 330 mM NaCl, 25 U/mL Benzonase, and protease inhibitor cocktail) and incubated for 5 minutes on ice. The tubes were mixed and then incubated for another 30 minutes on ice, then sonicated 5 times for 10 seconds each, with at least 30 seconds on ice between each round of sonication. Both nuclear and cytoplasmic samples were then centrifuged at 4 °C for 20 minutes at 20,000 g, and the pellet was discarded. 4X LDS sample buffer (Thermo NP0007) and BME were added to the supernatants to a final concentration of 1X and 1%, respectively, then heated to 95 °C for 5-10 minutes before loading on a gel.

The protein samples were run on a 7.5% (Biorad 4561024) or 8-16% gel (Thermo Scientific XP08160BOX), then transferred to a nitrocellulose membrane (GE Life Sciences #10600002). Membranes were incubated in blocking buffer (Thermo Scientific #37539), followed by the specified primary and then secondary antibodies. All incubations were done either at room temperature for 1 hour, or at 4 °C overnight. Signal was developed using SuperSignal™ West Pico PLUS Chemiluminescent Substrate (Thermo Scientific #34580) and images were taken on a ChemiDoc™ imager. When detecting PALI1 or JARID2, SuperSignal™ West Atto Ultimate Sensitivity Substrate (Thermo A38554) was added to the Pico PLUS substrate to a final concentration of 20%. Densitometry of the bands was measured using BioRad Image Lab.

The antibodies used for immunoblotting included: anti-GAPDH (Proteintech 10494-1-AP, 1:4000), anti-H3 (Abcam #Ab1791, 1:100000), anti-H3K27me1 (Active Motif 61015, 1:2000), anti-H3K27me2 (Abcam ab24684, 1:1000), anti-H3K27me3 (Active Motif 61017, 1:2500), anti-EZH2 (Cell Signalling 5246S, 1:1000), anti-SUZ12 (Cell Signalling 3737, 1:1000), anti-EED (Cell Signalling 85322, 1:1000), anti-LCOR (Merck #ABE1367, 1:250), anti-JARID2 (Cell Signalling 13594S, 1:1000), anti-LAMINB1 (Abcam ab16048, 1:10000), anti-mouse-HRP-conjugated (Jackson Immuno-Reasearch #715-035-150, 1:5000), and anti-rabbit HRP-conjugated (Santa Cruz Biotechnology #sc-2357, 1:5000).

### Immunofluorescence Confocal and STED Microscopy

Approximately 3–4 × 10⁴ cells were cytospun onto glass slides and allowed to air-dry slightly before fixation in 4% paraformaldehyde solution (PFA) in PBS (Santa Cruz sc-281692) for 15 min at room temperature. Slides were washed three times with PBS, permeabilized with 0.1% Tween 20 in PBS for 10 min, and blocked in 1% BSA in PBS for 30 min. Primary antibodies against SUZ12 (Cell Signalling, 3737S), JARID2 (Cell Signalling, 13594S), RING1B (Cell Signalling, 5694S), and H3K27me3 (Active Motif, 61017) were diluted 1:100 in blocking buffer, whereas anti-PALI1 (Merck, ABE1367) antibody was diluted 1:50. Slides were incubated overnight at 4 °C in a humidified chamber, washed three times with PBS, and incubated for 1 h at room temperature with fluorophore-conjugated secondary antibodies (Alexa 488 anti-mouse (Thermo, A-11001) and Alexa 647 anti-rabbit (Thermo, A32795) for confocal imaging, or Alexa 594 anti-mouse (Thermo, AB_2535789) and STAR 635P anti-rabbit (Abberior, ST635P-1002-500UG) for STED; all at 1:500). For STED imaging, nuclei were counterstained with SYBR Green I (Thermo, S7563, 1:20,000) to visualize DNA, whereas for confocal imaging, nuclei were stained with DAPI (Sigma, D9542-1MG, 0.5 µg mL⁻¹) for 10 min. Slides were post-fixed in 4% PFA in PBS for 10 min, washed, air-dried, and mounted in Vectashield medium.

Confocal imaging was performed on a Zeiss LSM 980 microscope equipped with an Airyscan 2 detector and a 63×/1.4 NA oil-immersion objective. Images were acquired with a 5× zoom, pixel size 0.04 µm, frame size 763 × 763 pixels, and 2.0× sampling in SR mode. Frame time was 45.1 s (scan speed 9, maximum). Scans were unidirectional, with 16× line averaging, repeat-per-line acquisition, mean-intensity mode, and 8-bit depth. Excitation lasers were 405 nm (0.5%) for DAPI, 488 nm (6.5%) for Alexa 488, and 639 nm (11%) for Alexa 647, each with master gain 749 V and digital gain 1.0. Representative confocal localization images were obtained from at least three independent experiments, with ≥5 fields per experiment per sample acquired under identical settings.

Imaging was also performed on an Abberior Expert Line STED system (Olympus IX83 stand) equipped with a 100×/1.4 NA UPL SAPO oil-immersion objective and pulsed lasers at 488, 560, 595, 640, and 775 nm. Both confocal-mode and STED-mode images were acquired from the same fields using an identical pixel size of 30 nm, with images collected at a 5 µs pixel dwell time. The pinhole diameter was 1.0 Airy unit. In confocal mode, the depletion laser was off. Excitation powers were 12% for STAR 635P, 40% for STAR 580 (Alexa 594 equivalent), and 5% for STAR Green (SYBR Green I), each acquired with a single frame accumulation, except STAR 580, which was acquired with three frame accumulations. In STED mode, the same fields were sequentially acquired with the depletion laser on. Excitation/depletion powers were 24.1% excitation / 15% depletion for STAR 635P and 55.6% excitation / 30% depletion for STAR 580. STED images were collected with three-frame accumulation and 8-bit depth, achieving a lateral resolution of approximately 120–150 nm.

Representative STED images were obtained from one independent experiment, with 5–6 fields per sample captured under identical parameters. Representative images were processed in Fiji (ImageJ) for visualization. For quantitative analysis, STED imaging was performed in three independent experiments using identical acquisition settings. Approximately 15 nuclei per condition were initially imaged, and an additional 5–10 nuclei were acquired to compensate for segmentation exclusions. After quality control, 13–20 nuclei per condition per experiment were included for analysis. Negative-control samples were acquired as two images per experiment under identical conditions. Foci number and size were quantified using ilastik and CellProfiler, and data were analyzed at the experiment level.

### STED Image Processing and Quantification

Files generated by the Abberior STED system were batch-processed using a custom macro in Fiji/ImageJ 2.16.0/1.54p ^74^. This macro automatically converted the 16-bit .obf files into 8-bit single-channel .tif format. STED images captured in the 635-nm far-red channel, corresponding to RING1B, were automatically copied and saved into a separate directory for downstream analysis.

RING1B puncta were segmented using the interactive machine-learning software ilastik 1.4.0 (OSX) ^75^ with a set of classifiers specifically trained for these datasets. The simple segmentation masks output from the Pixel Classification workflow were converted from .hdf5 to .tif using ilastik’s built-in conversion function. Because these files are saved as unsigned integer format, they were not accurately interpreted by CellProfiler, necessitating an additional custom ImageJ/Fiji macro that automatically applied thresholding and converted the masks to 8-bit binary images (pixel intensities 0 or 255).

The binary masks and other channels were imported into CellProfiler 4.2.6 (OSX) ^83,84^ and processed using a custom analysis pipeline. RING1B binary masks and the nuclear channel (SYBR Green I) were segmented into individual objects using the IdentifyPrimaryObjects module. For H3K27me3 signals, background noise was first reduced using ReduceNoise (Size = 5, Distance = 1, Cutoff Distance = 0.1), followed by segmentation via IdentifyPrimaryObjects.

Each step of the pipeline was validated against negative-control samples, in which primary antibodies were omitted, to confirm segmentation accuracy. To ensure that each image contained a single complete nucleus, nuclei touching image borders were excluded. Using the RelateObjects module, H3K27me3 and RING1B objects belonging to the excluded nuclei were removed. The same module was applied to select RING1B objects overlapping with H3K27me3 signals, thereby identifying RING1B puncta associated with chromatin regions.

Morphometric features including area, MajorAxisLength, MeanRadius, and perimeter were extracted from total and H3K27me3-overlapping RING1B puncta using the MeasureObjectSizeShape module. Object counts and measurements were exported as .csv files via ExportToSpreadsheet. For segmentation quality control, object outlines were automatically overlaid onto the nuclear channel for each image, and these validation images were saved in the output directory.

All custom ImageJ macros, ilastik workflows, and CellProfiler pipelines used in this study are available at: https://github.com/ykim-codes/STED_Ring1b_ImageAnalysis. Data was deposited to Mendeley (DOI: 10.17632/3h6hxgrg2f.1).

### In vitro histone methyltransferase assays

Assays were performed using 100 nM PRC2 in 384 well microplates (Corning CLS4513) in a total reaction volume of 16 µL in reaction buffer (50 mM Tris pH 8.0 at 30 °C, 100 mM KCl, 0.5 mM MgCl2, 0.1 % Tween 20, 5 mM DTT, 25 µM SAM). Reactions were incubated at 30 °C for up to three hours as indicated in the text and quenched using 2 µl 0.5 % (v/v) trifluoroacetic acid. Luminescent signal was recorded using a MTase-Glo Methyltransferase Assay kit (Promega V7601), and measured using a CLARIOstar Plus Multi-Mode Plate Reader (BMG 8000-89). A SAH standard curve was generated for each independent replicate by plotting SAH against luminescence. SAH generated by PRC2 was interpolated from these standard curves. Three independent replicates were performed on separate days. The data points in each independent replicate represent the mean of two technical replicates after subtracting the mean of a negative control sample lacking mononucleosome substrate. Data was plotted and analysed using GraphPad Prism. The non-linear regression analysis for the Michaelis-Menten model of enzyme kinetics was applied to calculate *V_max_*, *K_M_*, and *k_cat_*.

The sequences of the peptides used in the assays are given below (a C-terminal Y was added to facilitate quantification):

H3K27me3: SKAARK(me3)SAPSTY

JARID2 K116me3: RLQAQRK(me3)FAQSQY

PALI1 K1241me3: KKHLKK(me3)FPGATY

### Quantitative chromatin immunoprecipitation with whole genome sequencing (ChIP-Rx)

Cells were collected by replacing the culture media with trypsin for <5 minutes, then resuspending the detached cells in growth media and centrifuging for 5 minutes at 300 g. The cells were first washed once in PBS, then crosslinked for 10 minutes at room temperature in PBS with 1% (v/v) formaldehyde. The formaldehyde was quenched by adding glycine to a final concentration of 0.125 M, and was then washed twice in PBS. Cell pellets containing an equal number of cells in each sample were snap frozen with liquid nitrogen and stored at -80°C.

The cell pellets were thawed on ice and lysed in lysis buffer (100 mM NaCl, 0.5% (v/v) SDS, 5 mM EDTA, 50 mM Tris-HCl pH 8 and 1X protease inhibitor cocktail (Sigma #4693132001)) on ice, then centrifuged for 6 minutes at 300 g. The resulting chromatin pellet was resuspended in ChIP buffer (a 2:1 ratio of lysis buffer and dilution buffer (100 mM NaCl, 5% (v/v), Triton X-100, 5mM EDTA, 100mM Tris-HCl pH 8.5 and 1X protease inhibitor cocktail (Sigma #4693132001))). For every 1×10^7^ cells, chromatin from 5×10^6^ cross-linked NTERA-2 cells were added. All samples were then sonicated at 4°C in a Bioruptor set to high, with 30 cycles of 30 s on and 30 s off. The sonicated samples were diluted to 1×10^7^ cells per mL, and 1mL per antibody was used for ChIP, with 1% kept as input.

Antibodies were added to each sample as specified and incubated overnight at 4 °C rotating at low speed. The antibodies used for ChIP included: anti-SUZ12 (Cell Signalling 3737, 0.42 µg), anti-H3K27me3 (Cell Signalling 35861SF, 3 µg) anti-Jarid2 (Cell Signalling 13594S, 1.115 µg), anti-MTF2 (Proteintech 16208-1-AP, 1.435 µg), anti-H3K4me3 (Cell Signalling 9751S, 0.605 µg), anti-H2AK119ub (Cell Signalling 8240S, 2.69 µg), and anti-RING1B (Cell Signalling 5694S, 1.095 µg). After the antibody incubation, the lysate was centrifuged at max speed for 30 minutes at 4 °C. The pellet was discarded and 40 µL of Dynabeads Protein G (Thermo #10003D), washed 5 times in ChIP buffer, were added to the supernatant of each sample and incubated for 2-4 hours at 4°C rotating at low speed. The beads were then washed 3 times in mixed micelle buffer (150 mM NaCl, 20 mM Tris-HCl pH 8, 5 mM EDTA, 5.2% (w/v) sucrose and 1% (v/v) Triton X-100), twice in buffer 500 (0.1% (w/v) sodium deoxycholate, 1 mM EDTA, 50 mM HEPES pH 7.5 and 1% (v/v) Triton X-100), twice in LiCl/detergent wash buffer (0.5% (w/v) sodium deoxycholate, 1 mM EDTA, 250 mM LiCl, 0.5% (v/v) NP-40, 10 mM Tris-HCl pH 8) and once in TE (10 mM Tris-HCl pH 8 and 1 mM EDTA). The beads were then resuspended in 100 µL elution buffer (1% (v/v) SDS and 100 mM NaHCO3) and incubated at 65°C for 1 hour with shaking. The eluate was removed from the beads, and 10 µL of the input was combined with 90 µL of elution buffer, then all samples were reverse-crosslinked by shaking at 65 °C overnight.

1 µL of RNase A (Thermo #EN0531) was then added to all samples and incubated shaking at 37 °C for 1 hour, followed by 1 µL of Proteinase K (NEB #P8107S) with 2 hours of shaking at 55 °C. DNA was purified from these samples using the Qiagen MinElute PCR purification kit (Qiagen #28006) according to the instructions of the manufacturer. The purified DNA was dried overnight at room temperature in a biosafety hood before shipment to Azenta. Library preparation and Illumina sequencing with 20 million paired-end reads of 150bp per library was performed by Azenta for all samples.

### Bioinformatic analysis of ChIP-Rx datasets

Chromosome names for the mouse genome were modified with the prefix ‘mm10_’ and a metagenome was created by concatenating the mouse and human (spikein) reference genomes (mm10 and hg38, respectively) before indexing with Bowtie2 ^80^ as described previously ^47^. Reads were aligned to the metagenome using Bowtie2 with default parameters ^80^. Non-unique read alignments were filtered out using SAMtools ^81^ to exclude those with an alignment quality of <2. The mm10_ prefix appended to chromosome names was used to separate reads aligned to the reference mouse or spike-in human genomes. SAMtools was used to convert SAM files to BAM files and to remove duplicate aligned reads. Spike-in normalization factors were calculated for each ChIP dataset using the formula for normalized reference-adjusted reads per million (RRPM) as described ^47^ (1 per million spike-in reads). BigWig files were generated using the bamCoverage tool from the deepTools suite (version 3.3.0) ^82^ with a bin size of 10 and data were subsequently visualized as ChIP-Rx normalized tracks using the IGV genome browser.

Transcription start sites (TSS) for the mm10 mouse genome build were defined and annotated as all unique protein coding TSS +/- 5kb (n=19,888; downloaded from ENSEMBL biomart). Peaks were called for SUZ12 from in the wild-type cells using MACS2 with paired-end function (BAMPE) and an FDR < 0.01 ^83^. mm10 blacklisted regions were removed from all peak sets prior to downstream analyses ^84^. “PRC2-targets” (n=4,445) were defined as TSS that overlapped with a SUZ12 peak using bedTools intersect with -v and -wa flags ^85^. The remaining TSS were defined as “non-PRC2 targets” (n=16,118). “Intergenic regions” (n=22,050) included all non-blacklist regions after excluding annotated genes and 5 kb upstream and downstream from the start (TSS) and end (TES) of annotated genes.The genome-wide Log_2_ fold-change of the ChIP-Rx signals for H3K27me3 and SUZ12 in the mutant cell line was calculated using the bigwigCompare tool from the deepTools suite ^82^. “Outside SUZ12 peak” regions were defined as regions that reside within 30 kb upstream or downstream from the boundaries of all SUZ12 peaks and do not otherwise overlap with other SUZ12 peaks. Reads from both inside and outside SUZ12 peak regions were quantified using featureCounts ^86^, Rx-normalised, and visualised as violin plots using the ggpubr R package. Pairwise statistical tests were also performed using the ggpubr package. The proportion of H3K27me3 reads overlapping with H3K27me3 peaks was calculated using featureCounts. For quartile analysis, wild type H3K27me3 broad peaks were called using MACS2 with paired-end function (BAMPE) and an FDR 1×10^-4 83^. The resulting 7,879 H3K27me3 peaks were then sorted by size and split into quartiles (top 25%, 25-50%, 50-75%, 75-100%) based on peak width. All heatmaps and average enrichment profiles were generated using the computeMatrix, plotHeatmap and plotProfile tools from the deepTools suite ^82^.

## References

1. Kim, J. J. & Kingston, R. E. Context-specific Polycomb mechanisms in development. Nat Rev Genet 23, 680–695 (2022).

2. Blackledge, N. P. & Klose, R. J. The molecular principles of gene regulation by Polycomb repressive complexes. Nature Reviews Molecular Cell Biology 22, 815–833 (2021).

3. Piunti, A. & Shilatifard, A. The roles of Polycomb repressive complexes in mammalian development and cancer. Nature Reviews Molecular Cell Biology 22, 326–345 (2021).

4. Faust, C., Schumacher, A., Holdener, B. & Magnuson, T. The eed mutation disrupts anterior mesoderm production in mice. Development 121, 273–85 (1995).

5. O’Carroll, D. et al. The polycomb-group gene Ezh2 is required for early mouse development. Mol Cell Biol 21, 4330–4336 (2001).

6. Pasini, D., Bracken, A. P., Jensen, M. R., Lazzerini Denchi, E. & Helin, K. Suz12 is essential for mouse development and for EZH2 histone methyltransferase activity. EMBO J 23, 4061–4071 (2004).

7. Pengelly, A. R., Copur, Ö., Jäckle, H., Herzig, A. & Müller, J. A histone mutant reproduces the phenotype caused by loss of histone-modifying factor Polycomb. Science 339, 698–699 (2013).

8. Condemi, L., Mocavini, I., Aranda, S. & Di Croce, L. Polycomb function in early mouse development. Cell Death Differ 1–10 (2024) doi:10.1038/s41418-024-01340-3.

9. Uckelmann, M. & Davidovich, C. Not just a writer: PRC2 as a chromatin reader. Biochemical Society Transactions 49, 1159–1170 (2021).

10. Grewal, S. I. S. The molecular basis of heterochromatin assembly and epigenetic inheritance. Mol Cell 83, 1767–1785 (2023).

11. Hansen, K. H. et al. A model for transmission of the H3K27me3 epigenetic mark. Nat Cell Biol 10, 1291–1300 (2008).

12. Margueron, R. et al. Role of the polycomb protein EED in the propagation of repressive histone marks. Nature 461, 762–767 (2009).

13. Qi, W. et al. An allosteric PRC2 inhibitor targeting the H3K27me3 binding pocket of EED. Nat Chem Biol 13, 381–388 (2017).

14. Lee, C. H. et al. Allosteric Activation Dictates PRC2 Activity Independent of Its Recruitment to Chromatin. Mol Cell 70, 422–434.e6 (2018).

15. Sanulli, S. et al. Jarid2 Methylation via the PRC2 Complex Regulates H3K27me3 Deposition during Cell Differentiation. Mol Cell 57, 769–783 (2015).

16. Zhang, Q. et al. PALI1 facilitates DNA and nucleosome binding by PRC2 and triggers an allosteric activation of catalysis. Nature Communications 12, 4592 (2021).

17. Hauri, S. et al. A High-Density Map for Navigating the Human Polycomb Complexome. Cell Rep 17, 583–595 (2016).

18. Conway, E. et al. A Family of Vertebrate-Specific Polycombs Encoded by the LCOR/LCORL Genes Balance PRC2 Subtype Activities. Mol Cell 70, 408–421.e8 (2018).

19. Ge, E. J., Jani, K. S., Diehl, K. L., Müller, M. M. & Muir, T. W. Nucleation and propagation of heterochromatin by the histone methyltransferase PRC2: geometric constraints and impact of the regulatory subunit JARID2. J Am Chem Soc 141, 15029–15039 (2019).

20. Peng, J. C. et al. Jarid2/Jumonji Coordinates Control of PRC2 Enzymatic Activity and Target Gene Occupancy in Pluripotent Cells. Cell 139, 1290–1302 (2009).

21. Shen, X. et al. Jumonji modulates polycomb activity and self-renewal versus differentiation of stem cells. Cell 139, 1303–14 (2009).

22. Li, G. et al. Jarid2 and PRC2, partners in regulating gene expression. Genes Dev 24, 368–80 (2010).

23. Kalb, R. et al. Histone H2A monoubiquitination promotes histone H3 methylation in Polycomb repression. Nat Struct Mol Biol 21, 569–571 (2014).

24. Yu, J. R., Lee, C. H., Oksuz, O., Stafford, J. M. & Reinberg, D. PRC2 is high maintenance. Genes Dev 33, 903–935 (2019).

25. Davidovich, C. & Zhang, Q. Allosteric regulation of histone lysine methyltransferases: from context-specific regulation to selective drugs. Biochemical Society Transactions 49, 591–607 (2021).

26. Yang, Y. & Li, G. Post-translational modifications of PRC2: signals directing its activity. Epigenetics & Chromatin 13, 47 (2020).

27. Park, S. H. et al. Going beyond Polycomb: EZH2 functions in prostate cancer. Oncogene 40, 5788–5798 (2021).

28. Liu, X. A Structural Perspective on Gene Repression by Polycomb Repressive Complex 2. Subcell Biochem 96, 519–562 (2021).

29. Hananya, N., Koren, S. & Muir, T. W. Interrogating epigenetic mechanisms with chemically customized chromatin. Nat Rev Genet 25, 255–271 (2024).

30. Chittock, E. C., Latwiel, S., Miller, T. C. R. & Müller, C. W. Molecular architecture of polycomb repressive complexes. Biochem Soc Trans 45, 193–205 (2017).

31. Kadoch, C., Copeland, R. A. & Keilhack, H. PRC2 and SWI/SNF Chromatin Remodeling Complexes in Health and Disease. Biochemistry 55, 1600–1614 (2016).

32. van Kruijsbergen, I., Hontelez, S. & Veenstra, G. J. C. Recruiting polycomb to chromatin. Int J Biochem Cell Biol 67, 177–187 (2015).

33. Holoch, D. & Margueron, R. Mechanisms Regulating PRC2 Recruitment and Enzymatic Activity. Trends Biochem Sci 42, 531–542 (2017).

34. Laugesen, A. & Helin, K. Chromatin Repressive Complexes in Stem Cells, Development, and Cancer. Cell Stem Cell 14, 735–751 (2014).

35. Takeuchi, T. et al. Gene trap capture of a novel mouse gene, jumonji, required for neural tube formation. Genes Dev 9, 1211–22 (1995).

36. Blackledge, N. P. et al. Variant PRC1 complex-dependent H2A ubiquitylation drives PRC2 recruitment and polycomb domain formation. Cell 157, 1445–1459 (2014).

37. Li, X. et al. Mammalian polycomb-like Pcl2/Mtf2 is a novel regulatory component of PRC2 that can differentially modulate polycomb activity both at the Hox gene cluster and at Cdkn2a genes. Mol Cell Biol 31, 351–64 (2011).

38. Isono, K. et al. Mammalian polyhomeotic homologues Phc2 and Phc1 act in synergy to mediate polycomb repression of Hox genes. Mol Cell Biol 25, 6694–706 (2005).

39. Suzuki, M. et al. Involvement of the Polycomb-group gene Ring1B in the specification of the anterior-posterior axis in mice. Development 129, 4171–4183 (2002).

40. Akasaka, T. et al. A role for mel-18, a Polycomb group-related vertebrate gene, during the anteroposterior specification of the axial skeleton. Development 122, 1513–1522 (1996).

41. van der Lugt, N. M. et al. Posterior transformation, neurological abnormalities, and severe hematopoietic defects in mice with a targeted deletion of the bmi-1 proto-oncogene. Genes Dev 8, 757–769 (1994).

42. Wang, S. et al. Polycomblike-2-deficient mice exhibit normal left–right asymmetry. Developmental Dynamics 236, 853–861 (2007).

43. Vizán, P. et al. The Polycomb-associated factor PHF19 controls hematopoietic stem cell state and differentiation. Science Advances 6, eabb2745 (2020).

44. Takihara, Y. et al. Targeted disruption of the mouse homologue of the Drosophila polyhomeotic gene leads to altered anteroposterior patterning and neural crest defects. Development 124, 3673–3682 (1997).

45. Schumacher, A., Faust, C. & Magnuson, T. Positional cloning of a global regulator of anterior–posterior patterning in mice. Nature 383, 250–253 (1996).

46. Pérez-Pérez, J. M., Candela, H. & Micol, J. L. Understanding synergy in genetic interactions. Trends Genet 25, 368–376 (2009).

47. Orlando, D. A. et al. Quantitative ChIP-Seq Normalization Reveals Global Modulation of the Epigenome. Cell Reports 9, 1163–1170 (2014).

48. Lavarone, E., Barbieri, C. M. & Pasini, D. Dissecting the role of H3K27 acetylation and methylation in PRC2 mediated control of cellular identity. Nat Commun 10, 1679 (2019).

49. Glancy, E. et al. PRC2.1- and PRC2.2-specific accessory proteins drive recruitment of different forms of canonical PRC1. Mol Cell 83, 1393–1411.e7 (2023).

50. Gail, E. H. et al. Inseparable RNA binding and chromatin modification activities of a nucleosome-interacting surface in EZH2. Nat Genet 56, 1193–1202 (2024).

51. Højfeldt, J. W. et al. Non-core Subunits of the PRC2 Complex Are Collectively Required for Its Target-Site Specificity. Mol Cell 76, 423–436.e3 (2019).

52. Healy, E. et al. PRC2.1 and PRC2.2 Synergize to Coordinate H3K27 Trimethylation. Mol Cell 76, 437–452.e6 (2019).

53. Kim, J.-A., Kwon, M. & Kim, J. Allosteric Regulation of Chromatin-Modifying Enzymes. Biochemistry 58, 15–23 (2019).

54. Iglesias, N. et al. Automethylation-induced conformational switch in Clr4 (Suv39h) maintains epigenetic stability. Nature 560, 504–508 (2018).

55. Tang, Q., Zhang, A., Sullivan, M., Toth, K. F. & Aravin, A. A. Auto-methylation of the histone methyltransferase SetDB1 at its histone-mimic motifs ensures the spreading and maintenance of heterochromatin. 2025.01.21.634156 Preprint at 10.1101/2025.01.21.634156 (2025).

56. Fong, K. et al. PALI1 promotes tumor growth through competitive recruitment of PRC2 to G9A-target chromatin for dual epigenetic silencing. Molecular Cell 82, 4611–4626.e7 (2022).

57. Schmitges, F. W. et al. Histone methylation by PRC2 is inhibited by active chromatin marks. Mol Cell 42, 330–341 (2011).

58. Cookis, T., Lydecker, A., Sauer, P., Kasinath, V. & Nogales, E. Structural basis for the inhibition of PRC2 by active transcription histone posttranslational modifications. Preprint at 10.1101/2024.02.09.579730 (2024).

59. van de Wetering, M. et al. Prospective derivation of a Living Organoid Biobank of colorectal cancer patients. Cell 161, 933–945 (2015).

60. Arai, E. et al. Multilayer-omics analysis of renal cell carcinoma, including the whole exome, methylome and transcriptome. Int J Cancer 135, 1330–1342 (2014).

61. Bonilla, X. et al. Genomic analysis identifies new drivers and progression pathways in skin basal cell carcinoma. Nat Genet 48, 398–406 (2016).

62. Hayward, N. K. et al. Whole-genome landscapes of major melanoma subtypes. Nature 545, 175–180 (2017).

63. Shain, A. H. et al. Exome sequencing of desmoplastic melanoma identifies recurrent NFKBIE promoter mutations and diverse activating mutations in the MAPK pathway. Nat Genet 47, 1194–1199 (2015).

64. Cosmic. COSMIC - Catalogue of Somatic Mutations in Cancer. https://cancer.sanger.ac.uk/cosmic.

65. Tate, J. G. et al. COSMIC: the Catalogue Of Somatic Mutations In Cancer. Nucleic Acids Research 47, D941–D947 (2019).

66. Chammas, P., Mocavini, I. & Di Croce, L. Engaging chromatin: PRC2 structure meets function. Br J Cancer 122, 315–328 (2020).

67. Morschhauser, F. et al. Tazemetostat for patients with relapsed or refractory follicular lymphoma: an open-label, single-arm, multicentre, phase 2 trial. Lancet Oncol 21, 1433–1442 (2020).

68. Motoyama, J., Kitajima, K., Kojima, M., Kondo, S. & Takeuchi, T. Organogenesis of the liver, thymus and spleen is affected in jumonji mutant mice. Mech Dev 66, 27–37 (1997).

69. Takeuchi, T., Kojima, M., Nakajima, K. & Kondo, S. jumonji gene is essential for the neurulation and cardiac development of mouse embryos with a C3H/He background. Mech Dev 86, 29–38 (1999).

70. Lee, Y. et al. Jumonji, a nuclear protein that is necessary for normal heart development. Circ Res 86, 932–8 (2000).

71. Rice, P., Longden, I. & Bleasby, A. EMBOSS: the European Molecular Biology Open Software Suite. Trends Genet 16, 276–277 (2000).

72. EMBOSS. http://emboss.open-bio.org/.

73. Chang, Y.-C. et al. Nr6a1 controls Hox expression dynamics and is a master regulator of vertebrate trunk development. Nature Communications 13, 7766 (2022).

74. Rigueur, D. & Lyons, K. M. Whole-mount skeletal staining. Methods Mol Biol 1130, 113–121 (2014).

75. Hauswirth, G. M. et al. Breaking constraint of mammalian axial formulae. Nat Commun 13, 243 (2022).

76. Salmon provides fast and bias-aware quantification of transcript expression | Nature Methods. https://www.nature.com/articles/nmeth.4197.

77. Love, M. I., Huber, W. & Anders, S. Moderated estimation of fold change and dispersion for RNA-seq data with DESeq2. Genome Biol 15, 550 (2014).

78. Blain, R. et al. A tridimensional atlas of the developing human head. Cell 186, 5910 (2023).

79. Young, L., Sung, J., Stacey, G. & Masters, J. R. Detection of Mycoplasma in cell cultures. Nat Protoc 5, 929–934 (2010).

80. Langmead, B. & Salzberg, S. L. Fast gapped-read alignment with Bowtie 2. Nat Methods 9, 357–359 (2012).

81. Li, H. et al. The Sequence Alignment/Map format and SAMtools. Bioinformatics 25, 2078–2079 (2009).

82. Ramírez, F. et al. deepTools2: a next generation web server for deep-sequencing data analysis. Nucleic Acids Res 44, W160–165 (2016).

83. Zhang, Y. et al. Model-based analysis of ChIP-Seq (MACS). Genome Biol 9, R137 (2008).

84. Amemiya, H. M., Kundaje, A. & Boyle, A. P. The ENCODE Blacklist: Identification of Problematic Regions of the Genome. Sci Rep 9, 9354 (2019).

85. Quinlan, A. R. & Hall, I. M. BEDTools: a flexible suite of utilities for comparing genomic features. Bioinformatics 26, 841–842 (2010).

86. Liao, Y., Smyth, G. K. & Shi, W. featureCounts: an efficient general purpose program for assigning sequence reads to genomic features. Bioinformatics 30, 923–930 (2014).

